# Cross kingdom analysis of putative quadruplex-forming sequences in fungal genomes: novel antifungal targets to ameliorate fungal pathogenicity?

**DOI:** 10.1101/2020.09.23.310581

**Authors:** Emily F. Warner, Natália Bohálová, Václav Brázda, Zoë A. E. Waller, Stefan Bidula

**Affiliations:** School of Clinical Medicine, University of Cambridge, Cambridge, United Kingdom; Institute of Biophysics of the Czech Academy of Sciences, Brno, Czech Republic; Department of Experimental Biology, Faculty of Science, Masaryk University, Brno, Czech Republic; UCL School of Pharmacy, 29-39 Brunswick Square, London, United Kingdom; School of Biological Sciences, University of East Anglia, Norwich, United Kingdom

**Keywords:** Fungi, G-quadruplexes, i-motifs, *in-silico*, virulence, drug resistance

## Abstract

Fungi contribute to upwards of 1.5 million human deaths annually, are involved in the spoilage of up to a third of food crops, and have a devastating effect on plant and animal biodiversity. Moreover, this already significant issue is exacerbated by a rise in antifungal resistance and a critical requirement for novel drug targets. Quadruplexes are four-stranded secondary structures in nucleic acids which can regulate processes such as transcription, translation, replication, and recombination. They are also found in genes linked to virulence in microbes, and quadruplex-binding ligands have been demonstrated to eliminate drug resistant pathogens. Using a computational approach, we identified putative quadruplex-forming sequences (PQS) in 1362 genomes across the fungal kingdom and explored their potential involvement in virulence, drug resistance, and pathogenicity. Here we present the largest analysis of PQS in fungi and identified significant heterogeneity of these sequences throughout phyla, genera, and species. Moreover, PQS were genetically conserved. Notably, loss of PQS in cryptococci and aspergilli was associated with pathogenicity. PQS in the clinically important pathogens *Aspergillus fumigatus, Cryptococcus neoformans*, and *Candida albicans* were located within genes (particularly coding regions), mRNA, repeat regions, mobile elements, tRNA, ncRNA, rRNA, and the centromere. Genes containing PQS in these organisms were found to be primarily associated with metabolism, nucleic acid binding, transporter activity, and protein modification. Finally, PQS were found in over 100 genes associated with virulence, drug resistance, or key biological processes in these pathogenic fungi and were found in genes which were highly upregulated during germination, hypoxia, oxidative stress, iron limitation, and in biofilms. Taken together, quadruplexes in fungi could present interesting novel targets to ameliorate fungal virulence and overcome drug resistance.

## Introduction

Compared to viruses and bacteria, fungi are underappreciated [1]. They are key contributors to the food and drink, biotechnology, and textile industries, whilst also being an important source of novel antimicrobial compounds [2-5]. However, >1.5 million deaths per year are attributed to fungi in humans globally, more than malaria and on par with tuberculosis [6, 7]. Fungi can also cause blindness, serious skin conditions, promote allergic responses, and have been shown to cause secondary infections in cystic fibrosis, tuberculosis, Human Immunodeficiency Virus (HIV), and recently, SARS-CoV-2 patients [8, 9]. Fungal plant pathogens also destroy around a third of crops annually; enough to feed 600 million people [1]. Whilst fungal infections of amphibians, bats, bees, animals, and trees have a huge impact on biodiversity [7]. Thus, fungi are an important asset, but they also pose a devastating global burden, and a deeper understanding of their biology is therefore essential.

The negative effects of fungi are exacerbated by a lack of antifungals and the emergence of multidrug resistant pathogens with intrinsic resistance such as *Candida auris* and *Lomentospora prolificans* [7, 10]. Current classes of antifungals include azoles (e.g. fluconazole), echinocandins (e.g. caspofungin), and polyenes (e.g. amphotericin B), but resistance to these antifungals is ever more prevalent and there are no vaccines [11]. Indeed, a recent meeting of the World Health Organization (WHO) Expert Group on Identifying Priority Fungal Pathogens highlighted azole resistant *A. fumigatus* as one such priority pathogen [12]. Therefore, we have an urgent requirement to identify potential novel antifungal targets.

G-quadruplexes (G4s) and i-motifs (iMs) are intriguing four-stranded (quadruplex) secondary structures in nucleic acids that are enriched in regulatory regions, particularly the promoters and telomeric regions of prokaryotic and eukaryotic genomes [13, 14]. However, they can be found throughout the genome [15]. G4s can be found within guanine-rich regions of DNA and RNA. Here, four guanine bases associate through Hoogsteen hydrogen bonding to form the basic unit of the G4, the G-tetrad [16]. These can then stack on top of each other to form the G4 structure itself (Figure 1A-C). These stacks of G-tetrads are connected by loops of mixed-sequence nucleotides and can form intramolecular or intermolecular associations [17, 18]. This structure is further stabilised by the presence of monovalent cations, especially potassium [19]. Moreover, the 5’- to 3’-directionality of the strands, glycosidic bonding in the G-tetrads, the cation present, and number of stacked G-tetrads contribute to the wide variation of observed G4 structures and topologies [13]. Conversely, iMs form within cytosine-rich regions of DNA and can typically be found on the complementary strand opposite a G4 [20]. Like G4s, they are four stranded structures but are composed of two intercalated hairpins which are stabilised by hemi protonated cytosine-cytosine^+^ (C·C+) base pairs (Figure 1D-F) [21]. Studies into iMs have been limited compared to G4s as it was thought they weren’t physiologically relevant based on them being most stable in slightly acidic conditions. However, they have since been shown to form under physiological conditions, including neutral pH and molecular crowding, and have recently been identified within the nuclei of human cells [21-24]

**Figure 1.**
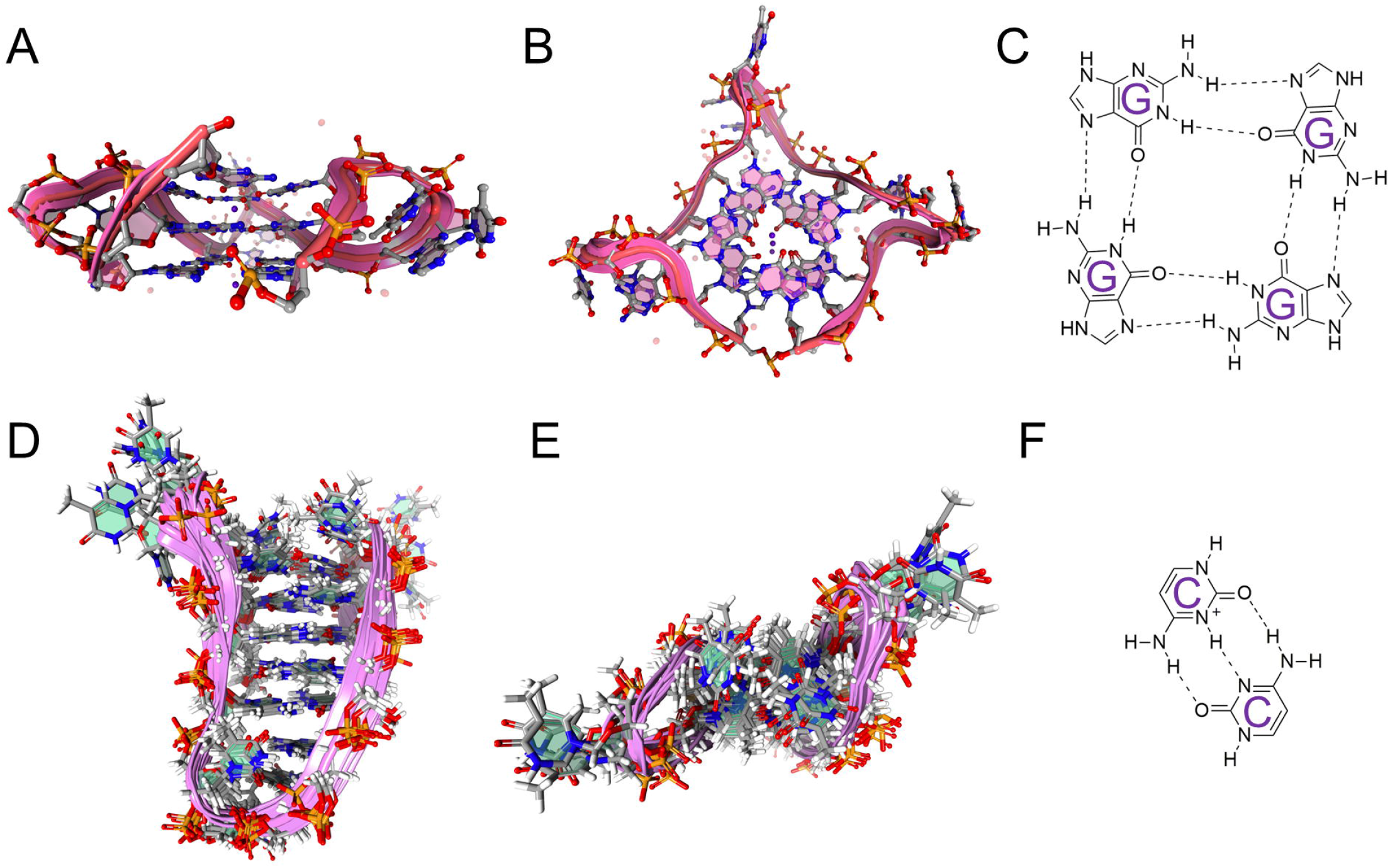
G-quadruplex (G4) and i-motif (iM) structures. Representative examples of G4s and iMs. **(A)** Side **(B)** and top-down view of the human telomere DNA quadruplex in K^+^ solution hybrid-1 form (PDB:2HY9). **(C)** The basic structure of the G-tetrad. **(D)** Side **(E)** and top-down view of an intramolecular i-motif DNA structure with C·C^+^ base pairing (PDB 1A83). **(F)** C·C^+^ base pairing found in i-motif structures. Images were generated using the Protein Imager software.

Once deemed structural curiosities, G4s and iMs have been highlighted to participate within host-pathogen interactions where they have important roles within gene regulation and have been shown to trigger phase separation of RNA in cells [25-27]. However, the regulation of biological functions by quadruplexes is complex and is influenced by their position within DNA/RNA, the surrounding topology, and environmental factors within the cell. In recent years, there has been increased interest in the therapeutic potential of targeting quadruplexes following the implication of these secondary structures in disease, especially cancer, due to their prevalence in oncogene promoters [26]. However, there is also now a growing number of pathogens in which G4s have been shown to contribute to virulence phenotypes, including; viruses (Human Papilloma Virus; Epstein-Barr Virus, HIV, SARS-CoV-2), prokaryotic bacterial pathogens (*Staphylococcus aureus, Streptococcus pneumoniae, Enterococcus* spp., *Borrelia* spp., *Neisseria meningitidis* and *N. gonorrhoeae*), and eukaryotic pathogens (*Trypanosoma brucei, Plasmodium falciparum*) [25, 28]. Notably, G4 DNA binding agents have been shown to be active against methicillin-resistant *S. aureus* and vancomycin-resistant *Enterococcus* spp.; potentially providing a novel target to overcome drug resistance [29, 30]. On the contrary, iMs have so far only been identified in the long terminal repeat promoter of the HIV-1 pro-viral genome where they modulate the transcription of viral genes [31]. Although, one could assume that this is only the tip of the iceberg, considering the ubiquitous nature of quadruplexes within the genomes of almost every organism.

Practically all organisms possess quadruplexes within their genomes but a thorough analysis of putative quadruplex-forming sequences (PQS) in fungi has not been conducted to date. Identifying PQS in key genes associated with virulence/drug resistance in fungal pathogens could highlight targets for G4/iM-binding ligands and a potential novel means to ameliorate fungal pathogenicity. In this study, we performed a computational investigation into the presence of PQS in 1362 genomes across the fungal kingdom and discussed their potential as novel antifungal targets within key pathogenic species.

## Methods

### Selection of genome sequences

The genomic sequences analysed in this study were obtained from publicly available repositories. The database (https://doi.org/10.5281/zenodo.3783970) was used for the Ascomycota [32]. Regarding the Ascomycota, publicly available genomes from a comprehensive study on Saccharomycotina were obtained, the NCBI’s Genome Browser was used to obtain basic information on the strains, and draft genomes from the subphyla Pezizomycotina and Taphrinomycotina were compiled from GenBank via FTP access number (ftp://ftp.ncbi.nlm.nih.gov/genomes/). Published genomes from the Basidiomycota, Mucoromycota, Zoopagomycota, Chytridiomycota, Microsporidia, and Cryptomycota were obtained from the Joint Genome Institute MycoCosm portal (https://mycocosm.jgi.doe.gov/mycocosm/home; last accessed 16/7/2020) [33]. Only the genomes from isolates with the highest assembly level and latest release date were included.

### The principle of the G4Hunter algorithm and process of analysis

The G4Hunter web application was used to identify PQS within fungal genomes [34]. The algorithm used in G4Hunter considers the G-richness and G-skewness of a genome. Each position within a sequence is given a score between −4 and 4, with G’s giving a positive score and C’s giving a negative score. A’s and T’s are neutral and have a score of 0. An increasing G4Hunter score (either positive or negative) correlates with increased propensity to form quadruplexes. A near-zero average score is indicative of a sequence that is most likely to form stable duplexes. The G4Hunter score is the arithmetic mean value of the sequence.

When analysing the genomes, the scored nucleic acid sequence was computed for a sliding window of 30 nucleotides and a threshold above 1.5 for stringent analyses, or a window of 25 nucleotides and thresholds between 1-2 for complete analyses. Regions in which the value of the mean score was above a threshold (either positive or negative) were extracted. Potential G4- or iM-forming sequences were all noted as PQS and were not treated differently.

To identify the location of PQS within annotated genomic features, the file containing the annotations for known genomic features within the genome of the selected fungi were downloaded from the NCBI database. The presence of PQS within a pre-defined genomic feature (e.g. gene, mRNA, mobile element) or within ±100 bp of these genomic features were analysed.

Analysis of the genomes for PQS was conducted using the G4Hunter DNA analyser web application (http://bioinformatics.ibp.cz:8888/#/) [34]. The location of PQS in known genomic features were identified using a publicly available script found at https://pypi.org/project/dna-analyser-ibp/. The gene ontologies and protein classes of genes containing PQS were determined via PANTHER™ GO slim v.15.0 and PANTHER™ Protein Class v.15.0, respectively [35, 36]. The sequences identified in G4Hunter were verified via QGRS Mapper (http://bioinformatics.ramapo.edu/QGRS/analyze.php) [37]. Sequences were analysed using the default settings of max length (30), min G-group (2) and loop size (0-36). The highest scoring sequences with the shortest loop length were selected in each case.

ChemDraw Ultra v12.0 was used to draw structures and The Protein Imager (https://3dproteinimaging.com/protein-imager/) [38] was used to generate 3D images from PDBs. These data were processed, and graphs were generated using GraphPad Prism software v6.01.

### Transcriptome analysis

Upregulated genes in germinating conidia and hyphae [39], during hypoxia [40], and during iron limitation or oxidative stress [41] were identified using publicly available transcriptome datasets. These were analysed in FungiDB (https://fungidb.org/) [42]. Upregulated genes were identified by comparing a reference sample (control) with a comparison sample (test). In each case, the difference between the minimum expression value of each gene in the reference sample and maximum expression value in the comparison sample was quantified. All genes upregulated >2-fold were acquired. Upregulated transcripts in *A. fumigatus* biofilms [43] were identified via the manuscript’s Supplementary Data. PQS in the top 20 most upregulated genes were identified using G4Hunter and QGRS Mapper using the default search settings.

### Statistics

Statistical analyses were conducted via parametric One-way ANOVA with Tukey’s multiple comparisons, Student’s t-test, or Pearson’s correlation coefficient; or non-parametric Kruskal-Wallis tests with post-hoc Dunn’s test and Bonferroni corrections using GraphPad Prism software v6.01. p<0.05 were considered statistically significant.

## Results

### There is large heterogeneity in the frequency and number of PQS across the fungal kingdom

A thorough analysis of putative quadruplex-forming sequences in fungi has not been conducted to date. Using G4Hunter, the number of PQS were quantified in 1362 fungal genomes of varying assembly, across 7 divisions, and 15 sub-divisions of fungi. Due to the high variability in genome size and chromosome number between fungal species, to normalise the data, the total number of PQS in addition to the frequency of PQS/kbp and number of PQS relative to the GC content (PQS/GC%) were noted.

Across the divisions, the Chytridiomycota had the largest average number of PQS (24,734 PQS) and highest PQS frequencies (0.555 PQS/kbp and 473 PQS/GC%; Figure 2). The Mucoromycota and Basidiomycota also had large numbers of PQS (19575 and 17108 PQS, respectively; Figure 2A and E). The Basidiomycota and Zoopagomycota had high PQS frequencies relative to genome size (0.445 and 0.373 PQS/kbp, respectively; Figure 2B and F). The Mucoromycota and Basidiomycota displayed high PQS frequencies relative to GC content (459 and 340 PQS/GC%, respectively; Figure 2C and G). Fungi within the Basidiomycota had the highest average GC content (53.3%; Figure 2D and H). The Microsporidia and Cryptomycota scored lowest for total number of PQS (300 and 372, respectively), PQS/kbp (0.091 and 0.029, respectively) and PQS/GC% (8 and 11, respectively; Figure 2). Moreover, they also had low GC content (39.6% and 35.0%, respectively).

**Figure 2.**
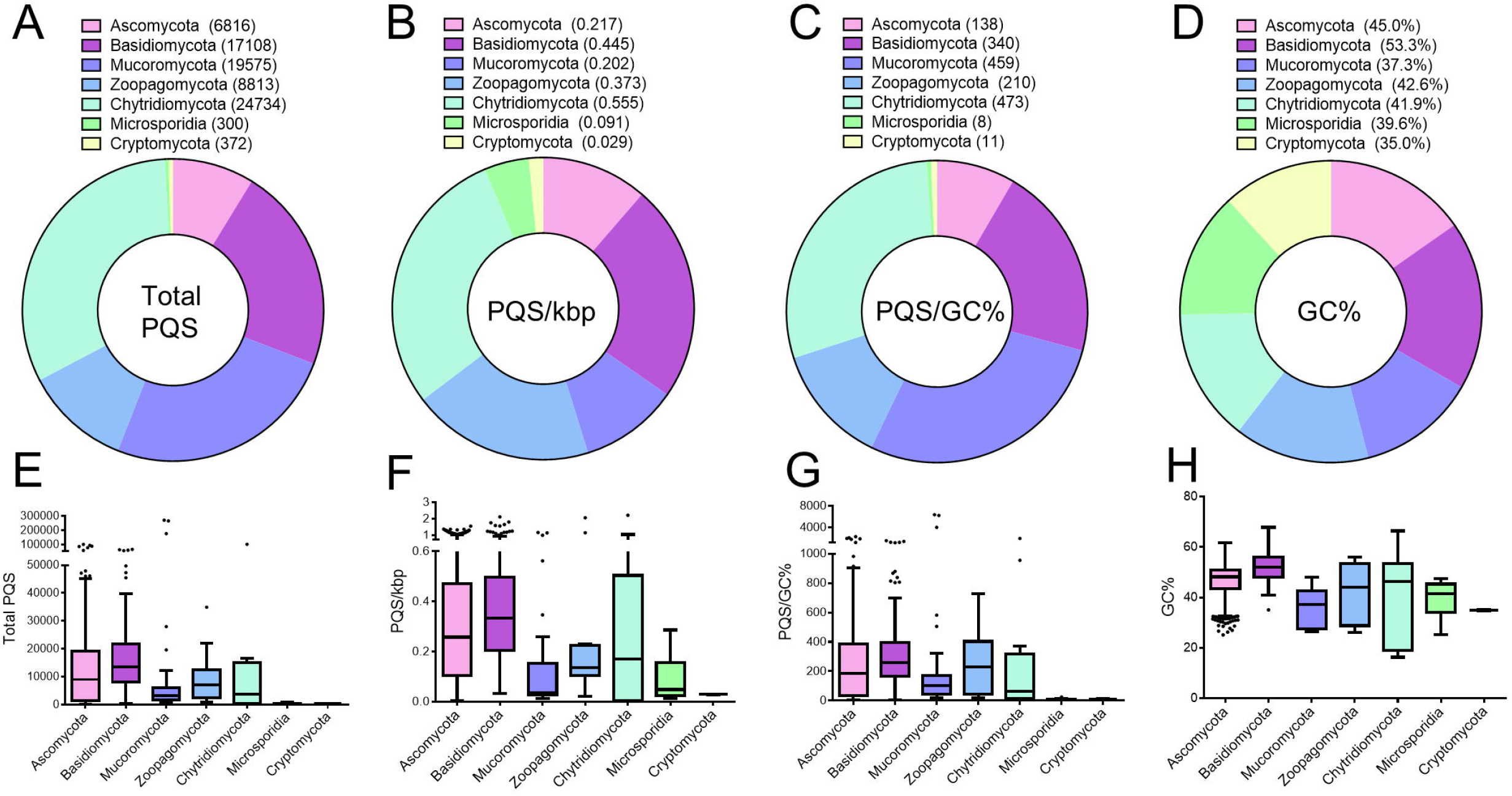
Heterogeneity of PQS across fungal divisions. The total number and frequency of PQS within 1362 fungal genomes were analysed using G4Hunter with a threshold of 1.5 and a window size of 30. The average number of PQS (**A** and **E**), PQS/kbp (**B** and **F**), PQS/GC% (**C** and **G**) and GC content (**D** and **H**) in fungi from the Ascomycota (n=1107), Basidiomycota (n=189), Mucoromycota (n=31), Zoopagomycota (n=12), Chytrdiomycota (n=12), Microsporidia (n=9), and Cryptomycota. (n=2). E to H contain boxplots with Tukey whiskers. The outliers are indicated by dots and the line within the boxplot is representative of the median value.

Considering G4s and iMs form in guanine or cytosine rich regions, respectively, one would expect fungi with a higher genome GC content to have a higher PQS frequency by chance. To investigate this further, the frequency of PQS/kbp relative to the GC content in all fungi and their divisions were plotted. As expected, there was a positive correlation between GC content and PQS frequency amongst all the fungal species analysed (r=0.5290; p<0.0001; Figure 3A). Moreover, this positive correlation was observed amongst the Ascomycota, Basidiomycota, and Mucoromycota (r=0.5619, r=0.3891, and r=0.5239 and r=0.2883, respectively; all p<0.05; Figure 3B-D). However, there was not a significant correlation observed within the Zoopagomycota, Chytridiomycota, Microsporidia, or Cryptomycota, possibly due to the limited sample size (Figure 3E-H).

**Figure 3.**
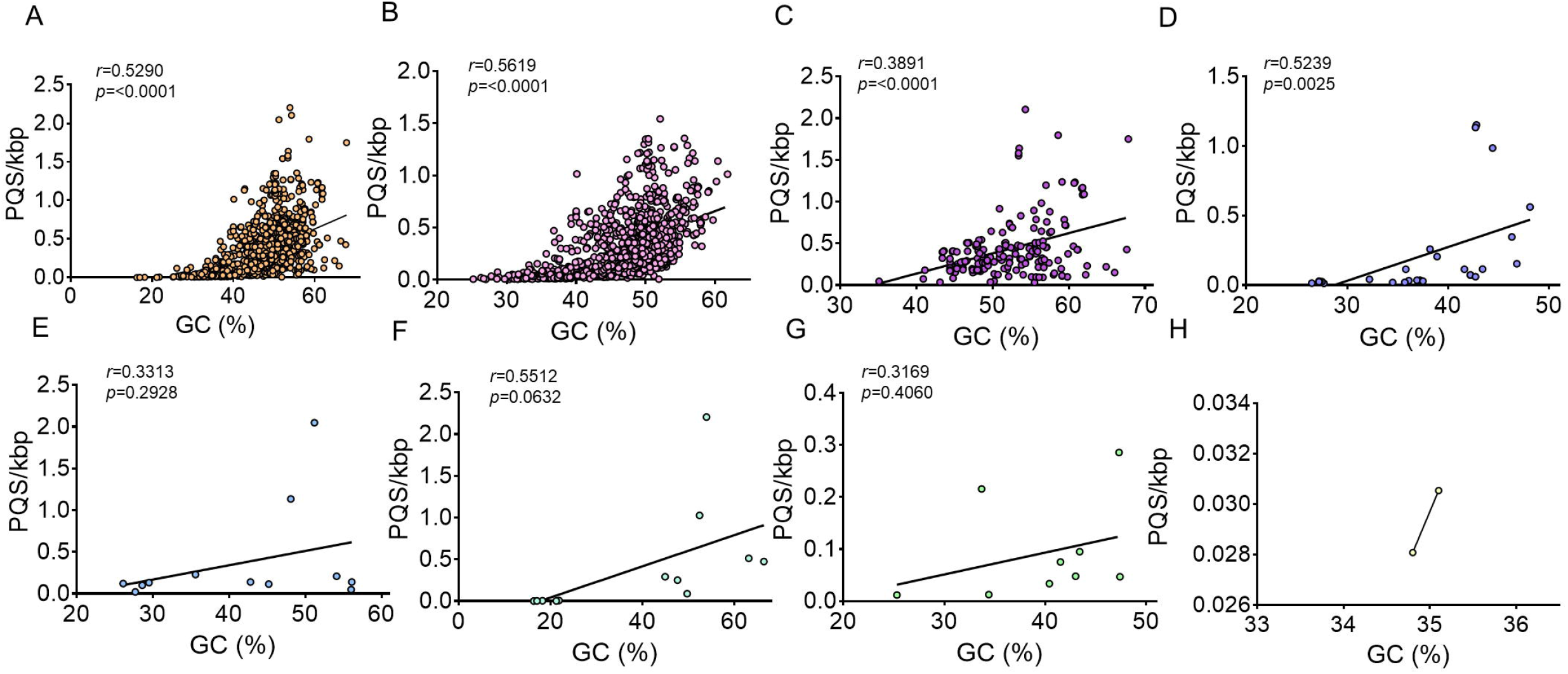
Higher genome GC-content is positively correlated with the frequency of PQS. The frequency of PQS relative to the GC content of fungi was plotted for **(A)** all fungal genomes in the study, **(B)** the Ascomycota, **(C)** the Basidiomycota, **(D)** the Mucoromycota, **(E)** the Zoopagomycota, **(F)** the Chytridiomycota, **(G)** the Microsporidia, and **(H)** the Cryptomycota. The Pearson correlation coefficient was used to determine the association between PQS and GC content. P<0.05 was considered statistically significant.

### There is large heterogeneity in the number and frequency of PQS within sub-divisions and in fungal genera containing important fungal pathogens

Within the Ascomycota, the average number of PQS in the sub-divisions Taphrinomycotina (n=14 species), Saccharomycotina (n=332 species), and Pezizomycotina (n=761 species) were 2799.7, 1079.0, and 17021.0, respectively (Figure 4A). The average PQS/kbp for each subphylum was 0.147, 0.075, and 0.436, respectively (Figure 4B). The average PQS/GC% for each subphylum was 42.4, 25.2, and 344.3, respectively (Figure 4C). Finally, the average GC% for each subphylum was 42.5%, 40.6%, and 49.6%, respectively (Figure 4D).

**Figure 4.**
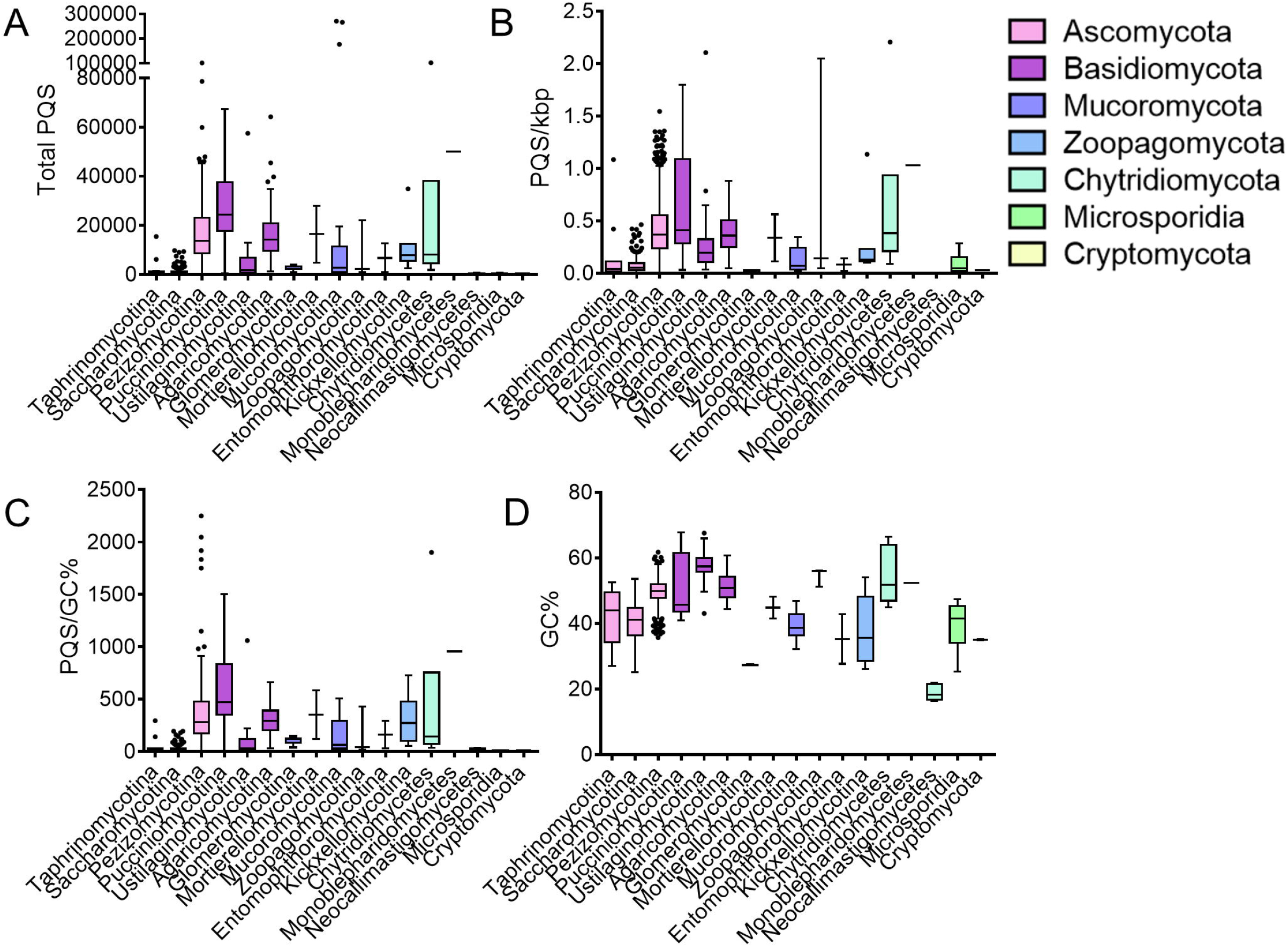
Heterogeneity of PQS across fungal sub-divisions. The total number and frequency of PQS within fungal sub-divisions were quantified using G4Hunter with a threshold of 1.5 and window size of 30. The average number of PQS **(A)**, PQS/kbp **(B)**, PQS/GC% **(C)** and GC content **(D)** in fungi from the Taphrinomycotina (n=14), Saccharomycotina (n=332), Pezizomycotina (n=761), Pucciniomycotina (n=24), Ustilagomycotina (n=31), Agaricomycotina (n=134), Glomeromycotina (n=9), Mortierellomycotina (n=2), Mucoromycotina (n=20), Zoopagomycotina (n=3), Entomophthoromycotina (n=2), Kickxellomycotina (n=7), Chytridiomycetes (n=6), Monoblepharidomycetes (n=1), Neocallimastigomycetes (n=5), Microsporidia (n=9), and Cryptomycota (n=2). A to D contain boxplots with Tukey whiskers. The outliers are indicated by dots and the line within the boxplot is representative of the median value.

Within the Basidiomycota, the average number of PQS in the sub-divisions Agaricomycotina (n=134 species), Pucciniomycotina (n=24 species), and Ustilagomycotina (n=31 species) were 16538.7, 29434.8, and 5351.0, respectively (Figure 4A). The average PQS/kbp for each subphylum was 0.433, 0.610, and 0.293, respectively (Figure 4B). The average PQS/GC% for each subphylum was 319.2, 605.6, and 94.1, respectively (Figure 4C). Finally, the average GC% for each subphylum was 51.1%, 51.1%, and 57.6%, respectively (Figure 4D).

Within the Mucoromycota, the average number of PQS in the sub-divisions Glomeromycotina (n=9 species), Mortierellomycotina (n=2 species), and Mucoromycotina (n=20 species) were 2900.1, 16444.0, and 39382.7, respectively (Figure 4A). The average PQS/kbp for each subphylum was 0.023, 0.338, and 0.246, respectively (Figure 4B). The average PQS/GC% for each subphylum was 106.5, 349.8, and 919.7, respectively (Figure 4C). Finally, the average GC% for each subphylum was 27.2%, 44.9%, and 39.7%, respectively (Figure 4D).

Within the Zoopagomycota, the average number of PQS in the sub-divisions Zoopagomycotina (n=3 species), Entomophthoromycotina (n=2 species), and Kickxellomycotina (n=7 species) were 8399.7, 6726.5, and 11315.1, respectively (Figure 4A). The average PQS/kbp for each subphylum was 0.746, 0.081, and 0.292, respectively (Figure 4B). The average PQS/GC% for each subphylum was 162.3, 162.7, and 305.4, respectively (Figure 4C). Finally, the average GC% for each subphylum was 54.4%, 35.3%, and 38.2%, respectively (Figure 4D).

Within the Chytridiomycota, the average number of PQS in the sub-divisions Chytridiomycetes (n=6 species), Monoblepharidomycetes (n=1 species), and Neocallimatigomycetes (n=5 species) were 23815.2, 50091, and 298, respectively (Figure 4A). The average PQS/kbp for each subphylum was 0.637, 1.027, and 0.003, respectively (Figure 4B). The average PQS/GC% for each subphylum was 446.7, 955.9, and 15.6, respectively (Figure 4C). Finally, the average GC% for each subphylum was 54.3%, 52.4%, and 18.9%, respectively (Figure 4D).

Within the Microsporidia (n=9 species) and Cryptomycota (n=2 species), the average number of PQS were 300.3 and 372, respectively (Figure 4A). The average PQS/kbp were 0.091 and 0.029, respectively (Figure 4B). The average PQS/GC% were 7.6 and 10.6, respectively (Figure 4C). Finally, the average GC% were 39.6% and 35.0%, respectively (Figure 4D).

Finally, we also highlighted the frequency of PQS in fungal genera which contained important human and plant pathogens. We found that there was also large heterogeneity in the frequency of PQS between species within genera containing human pathogens (e.g. *Aspergillus* spp., *Candida* spp., *Cryptococcus* spp., *Blastomyces* spp.) and plant pathogens (e.g. *Verticillium* spp., and *Fusarium* spp.; Figure 5). This variation was particularly wide within *Aspergillus* spp., and *Cryptococcus* spp.

**Figure 5.**
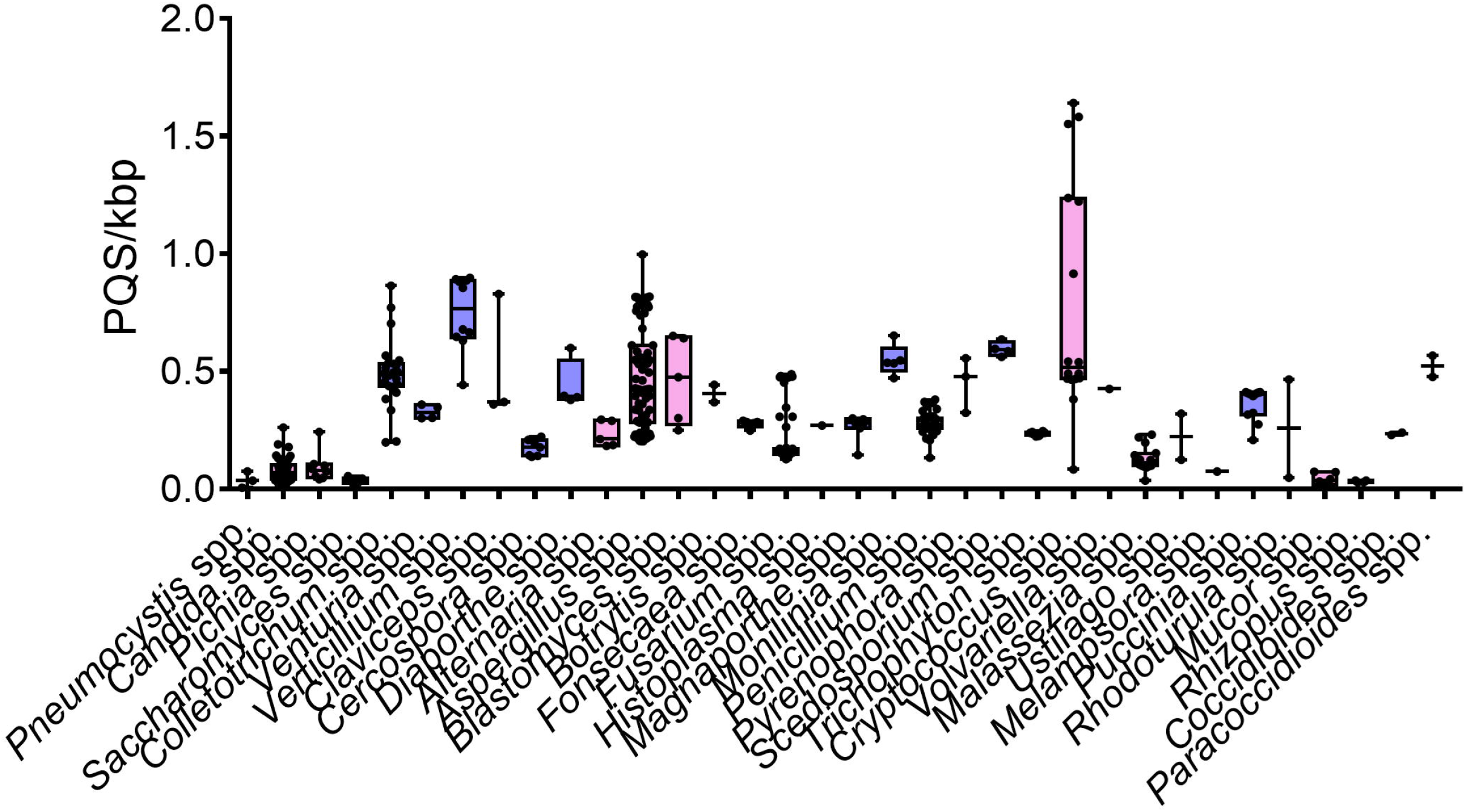
PQS distribution in fungal genera containing key species which cause disease in humans and plants. The frequency of PQS within genera containing key fungal pathogens were quantified using G4Hunter with a threshold of 1.5 and window size of 30. Fungal genera containing key plant pathogens are shown in blue, whilst genera containing key human pathogens are shown in pink. The boxplots display Tukey whiskers. The outliers are indicated by dots and the line within the boxplot is representative of the median value.

### PQS are evolutionarily conserved and are linked with genetic relatedness

Evolutionary conservation of genetic motifs within the genome are a hallmark of their fundamental importance to how that organism functions. Therefore, we endeavoured to explore whether there was evolutionary conservation of PQS within fungal genomes. We chose to explore this relationship in *Aspergillus* spp., due to the robustness and accuracy of the phylogenetic tree available [44].

Notably, we found that the frequency of PQS/kbp appeared to be intrinsically linked to how closely related species were, with species within the same section displaying similar PQS frequencies (Figure 6). Aspergilli in this tree were divided into 13 sections (range of PQS/kbp in brackets), the *Aspergillus* (0.364-0.461), the *Fumigati* (0.204-0.224), the *Candidi* (0.747-0.816), the *Circumdati* (0.395-0.467), the *Flavi* (0.225-0.289), the *Ochraceoros* (0.421-0.422), the *Usti* (0.385-0.396), the *Versicolores* (0.282-0.308), and the *Nigri* (0.495-0.782). The *Nigri* displayed the largest variation in PQS, however the lesser related species *A. carbonarius* and *A. aculeatus* skewed this range and if only the closest related species were noted, this range would be from 0.495-0.611 (Figure 6). As the *Nidulantes, Terrei, and Clavati* only contained one member each, correlations in these sections could not be made. Considering that we found a likely quadruplex-forming sequence within *cyp51A* in *A. fumigatus* and due to its importance in encoding a gene heavily involved in azole resistance, we chose to see if this sequence was conserved within the section *Fumigati*. Interestingly, *A. fischeri, A. novofumigatus*, and *A. lentulus* all retained the PQS observed in *A. fumigatus*, however this sequence could not be found in *A. udagawae* or *A. turcosus* (Figure 6).

**Figure 6.**
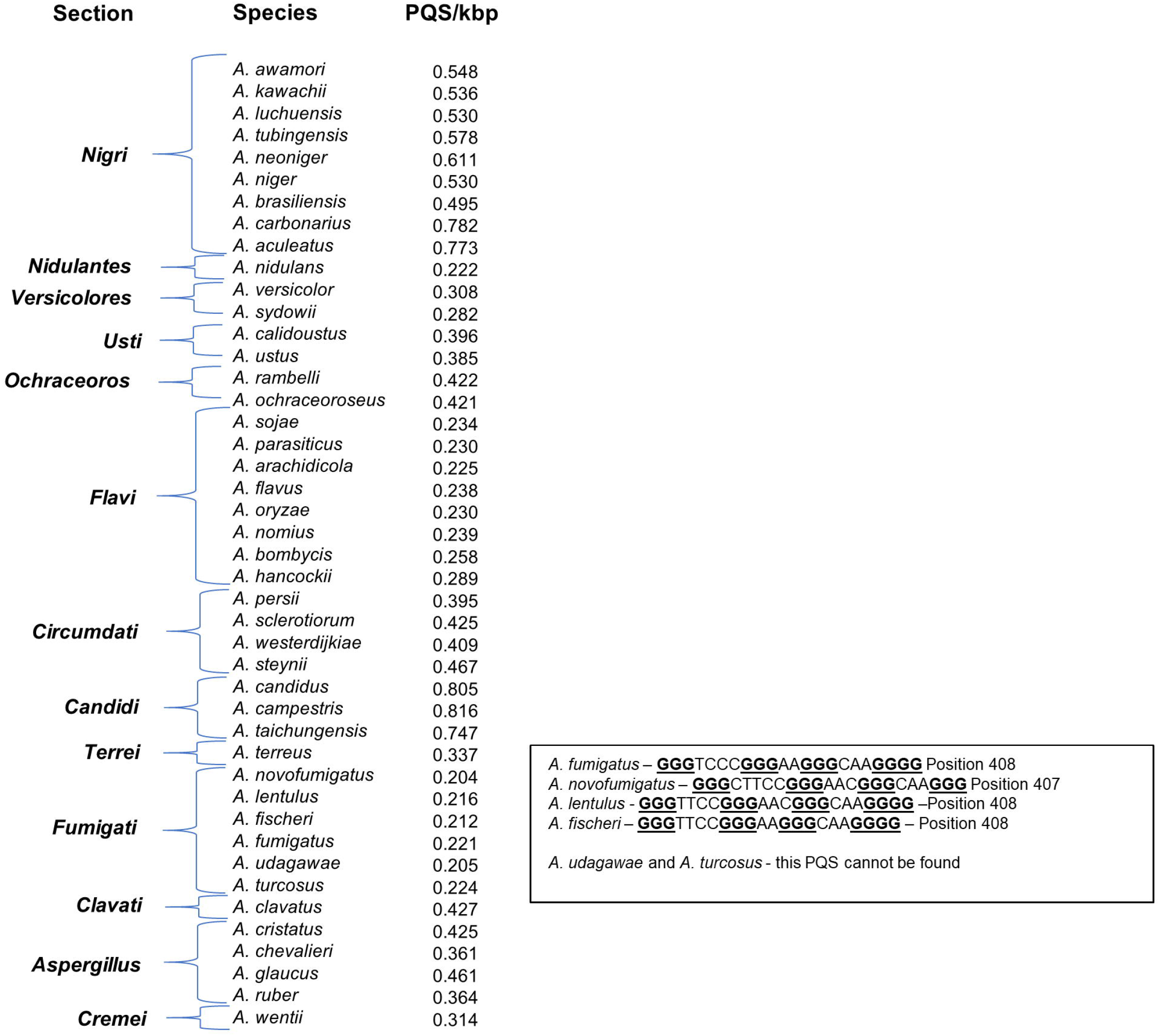
PQS in *Aspergillus* spp. are associated with genetic relatedness. *Aspergillus* spp., were categorised into sections based upon a phylogenetic tree generated by Steenwyk *et al*., [44]. The frequency of PQS was shown to be closely associated with the genetic relatedness of these fungi. Moreover, the PQS found in the *cyp51A* gene was conserved amongst species, but not found in *A. udagawae* or *A. turcosus*

### Loss of PQS is associated with pathogenicity in *Aspergillus* and *Cryptococcus* species, but not *Candida*

The Ascomycota and Basidiomycota contain many of the most prevalent fungal pathogens of both plants and humans, including the genera *Aspergillus* spp., *Candida* spp., and *Cryptococcus* spp., which contain fungal species that account for most fungal-related deaths in humans. Although, not all species within these genera are potential pathogens and we found high variation in their PQS frequency. Therefore, we compared the PQS frequency between pathogenic and non-pathogenic species to explore whether there was a link with pathogenicity.

Indeed, comparing 72 species of *Aspergillus* (22 pathogenic, 50 non-pathogenic) pathogenic species had a significantly lower frequency of PQS/kbp (0.364 vs. 0.523) and PQS/GC% (249.9 vs. 380.5) on average, compared to their non-pathogenic counterparts (Figure 7A and D). Further analysis highlighted that *A. unguis* had the lowest frequency of PQS/kbp (0.203 PQS/kbp), whilst *A. ellipticus* had the highest (0.997 PQS/kbp). Furthermore, the most pathogenic species, *A. fumigatus*, had a lower than average frequency of PQS (0.221 PQS/kbp; Supplementary Material). Notably, loss of PQS in the subphylum Pezizomycotina appeared to be associated with pathogenicity in humans overall (Supplementary Material).

**Figure 7.**
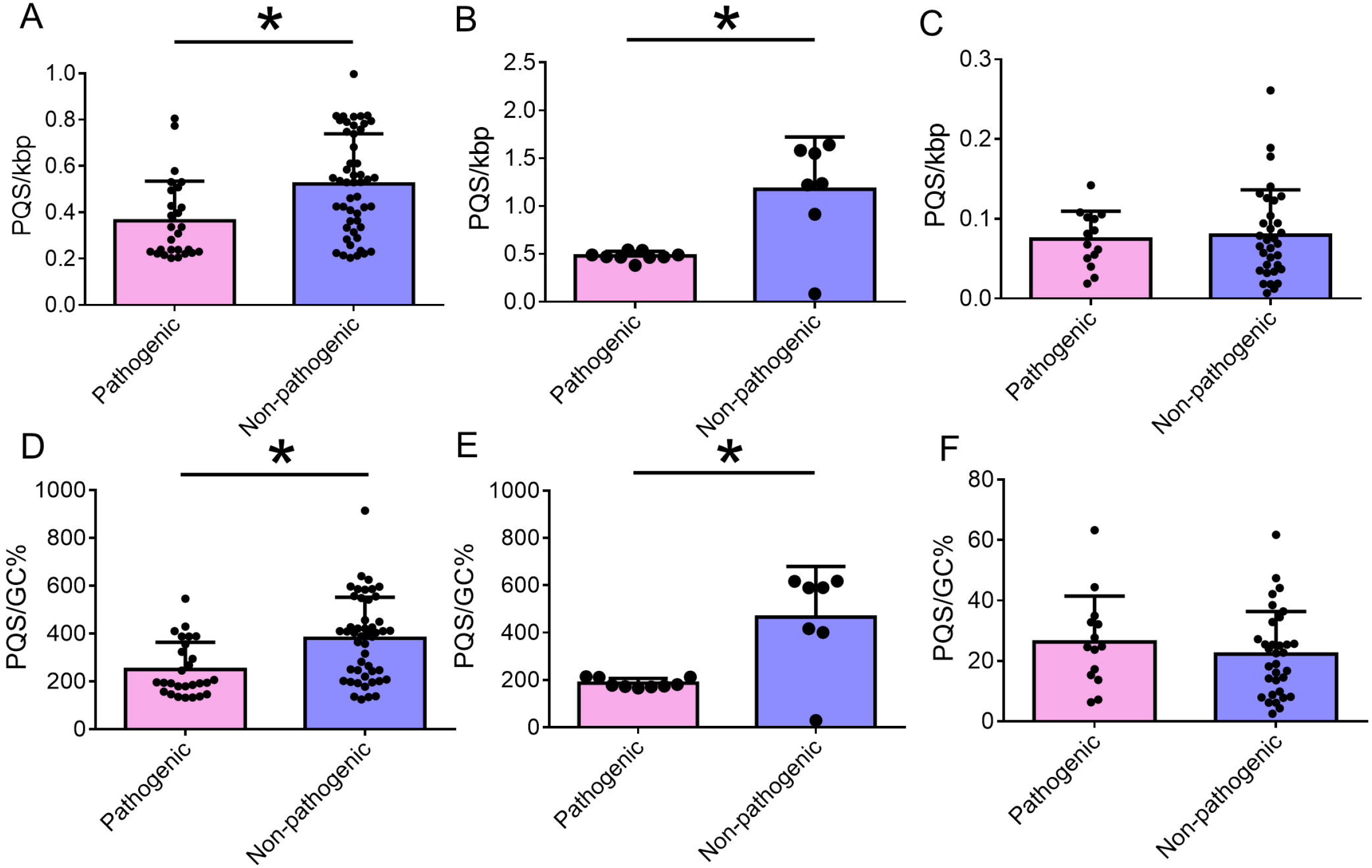
Pathogenic *Aspergillus* and *Cryptococcus* species have lower PQS frequencies compared to their non-pathogenic counterparts. The frequency of PQS/kbp and PQS/GC% were quantified and compared between pathogenic and non-pathogenic species within *Aspergillus* spp., *Cryptococcus* spp., and *Candida* spp. The PQS/kbp in pathogenic and non-pathogenic species of **(A)** *A. fumigatus*, **(B)** *C. neoformans*, and **(C)** *C. albicans*. The PQS/GC% in pathogenic and non-pathogenic species of **(D)** *A. fumigatus*, **(E)** *C. neoformans*, and **(F)** *C. albicans*. Dots represent individual species within a genus. The error bars represent the SD. Asterisks indicate p<0.05.

Similarly, comparing 16 species of *Cryptococcus* (9 pathogenic, 7 non-pathogenic) we also found that pathogenic species had a significantly lower frequency of PQS/kbp (0.480 vs. 1.176) and PQS/GC% (187.1 vs. 465.9) on average, compared to their non-pathogenic counterparts (Figure 7B and E). However, the lowest frequency of PQS/kbp was found in the non-pathogenic species *C. depauperatus* (0.084 PQS/kbp). Regarding pathogenic species, the *C. neoformans* AD hybrid was found to have the lowest PQS frequency (0.381 PQS/kbp). The highest PQS frequency was observed in *C. floricola* (1.64 PQS/kbp; Supplementary Material).

Conversely, this phenomenon was not observed in *Candida* spp., or closely related *Clavispora* and *Debaryomyces* species. Comparing 47 species (14 pathogenic, 33 non-pathogenic) we found that there was no association between PQS frequency and pathogenicity (Figure 7C and F). Pathogenic and non-pathogenic species had equivalent PQS/kbp (0.074 vs. 0.079) and PQS/GC% (26.3 vs. 22.3; Figure 7C and F). Moreover, this trend was also observed for pathogenic and non-pathogenic species across the entire subphylum Saccharomycotina (Supplementary Material). Further analysis showed that *C. orba* had the lowest frequency of PQS (0.006 PQS/kbp), whereas *C. fructus* had the highest (0.261 PQS/kbp; Supplementary Material). The most prevalent pathogenic species, *C. albicans*, had a higher than average frequency of PQS (0.101 PQS/kbp; Supplementary Material). Conversely, the multi-drug resistant species *C. auris* had both a lower frequency of PQS/kbp (0.061) and PQS/GC% (17.3), compared to the average.

### PQS are found in numerous genomic features in *A. fumigatus, C. neoformans*, and *C. albicans*

To evaluate the position of PQS within *A. fumigatus, C. neoformans*, and *C. albicans* (the most prevalent pathogens of their genera) the annotation information was obtained from the NCBI database and the presence of PQS within defined genomic features, or within 100 bp before or after these features were analysed. Both the total number of PQS per feature and the frequency of PQS/kbp relative to the combined genomic length of the described features were noted.

When only total PQS were considered, the largest number of PQS in all three fungal species could be found within the coding regions (CDS), genes, and mRNA, with few PQS found in other genomic features (Figure 8A, B, and C). However, this was not the same when considering the frequency of PQS/kbp of the genomic features. In *A. fumigatus*, the greatest frequency of PQS could be found in the repeat regions (Figure 8D). The lowest frequency could be found within the tRNA. In *C. neoformans*, the highest PQS frequencies were still in the CDS, genes, and mRNA, with a very low frequency found within the tRNA (Figure 8E). In *C. albicans*, the highest frequency of PQS could be found in the rRNA, followed by repeat regions and ncRNA (Figure 8F). There were no PQS found in the tRNA and low frequencies were again found in the mobile elements. The total number and frequency of PQS 100 bp before and after the annotated genomic features appeared to be evenly distributed (Figure 8).

**Figure 8.**
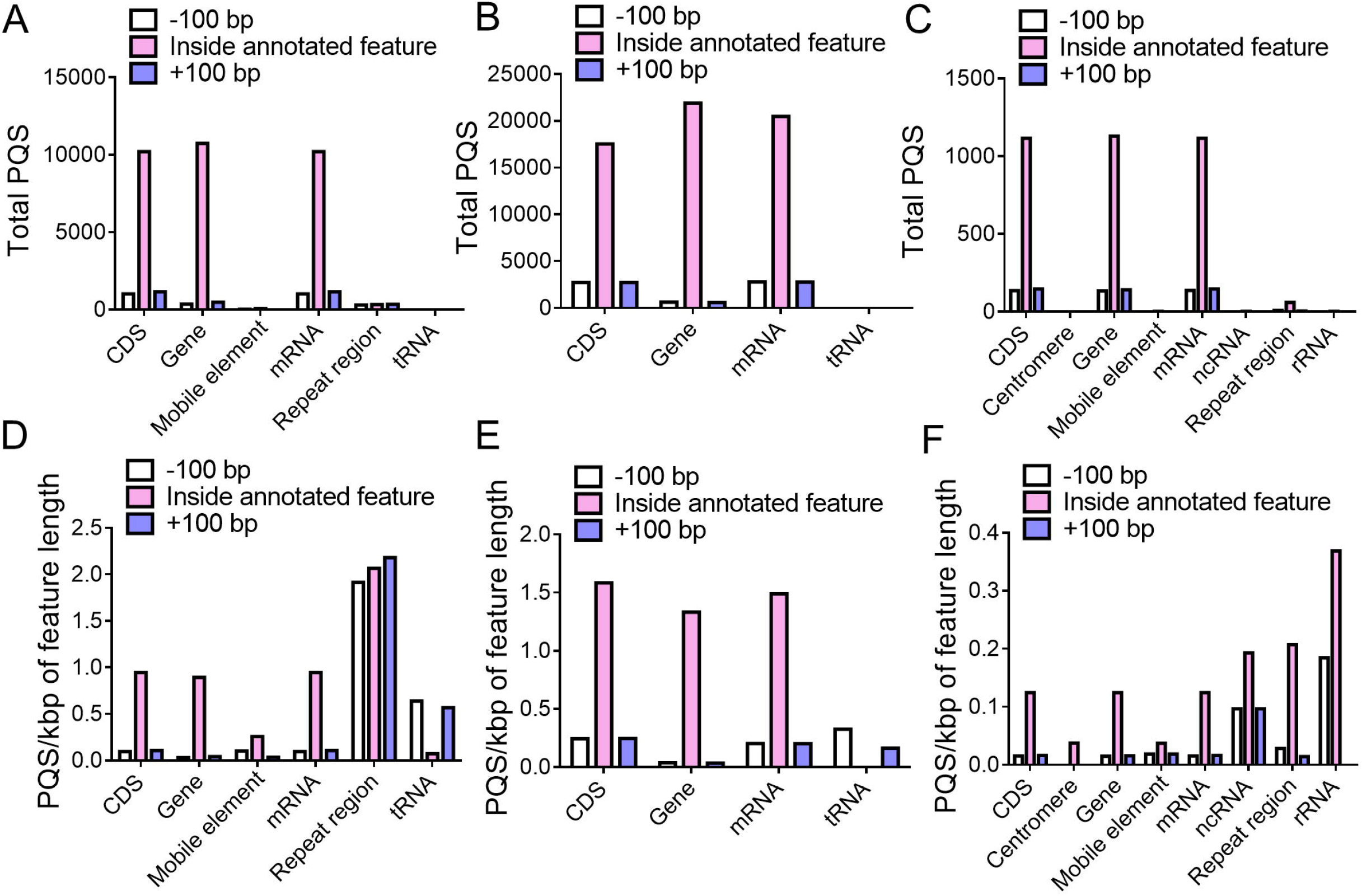
PQS in *A. fumigatus* Af293, *C. neoformans* JEC21, and *C. albicans* SC5314 can be found located throughout annotated genomic features. The location of PQS found 100 bp before, within, and 100 bp after annotated genomic features with a G4Hunter score >1.2. The total number of PQS in known genomic features in **(A)** *A. fumigatus*, **(B)** *C. neoformans*, and **(C)** *C. albicans*. The frequency of PQS comparative to the genomic length of the annotated features in **(D)** *A. fumigatus*, **(E)** *C. neoformans*, and **(F)** *C. albicans*.

### PQS are found in genes encoding proteins involved in metabolism, nucleic acid binding, cell transport, and protein modification

As we knew the genomic location of the PQS, we could then identify the number and identity of the genes which contained these sequences. This further enabled us to identify the classes of proteins associated with PQS-containing genes. In *A. fumigatus*, 35.1% of genes contained at least one PQS. In *C. neoformans*, this number was almost double, with 59.9% of genes containing PQS. Conversely, PQS were only found in 5.6% of genes in *C. albicans*.

Despite the discrepancies in the number of genes where PQS can found between the organisms, in all cases, PQS were primarily located in genes which encoded proteins involved in metabolism, nucleic acid binding, cell transport, and protein modification (Figure 9). They were least likely to be found in genes encoding for calcium-binding proteins, extracellular matrix proteins, cell adhesion molecules, and defense/immunity proteins (Figure 9).

**Figure 9.**
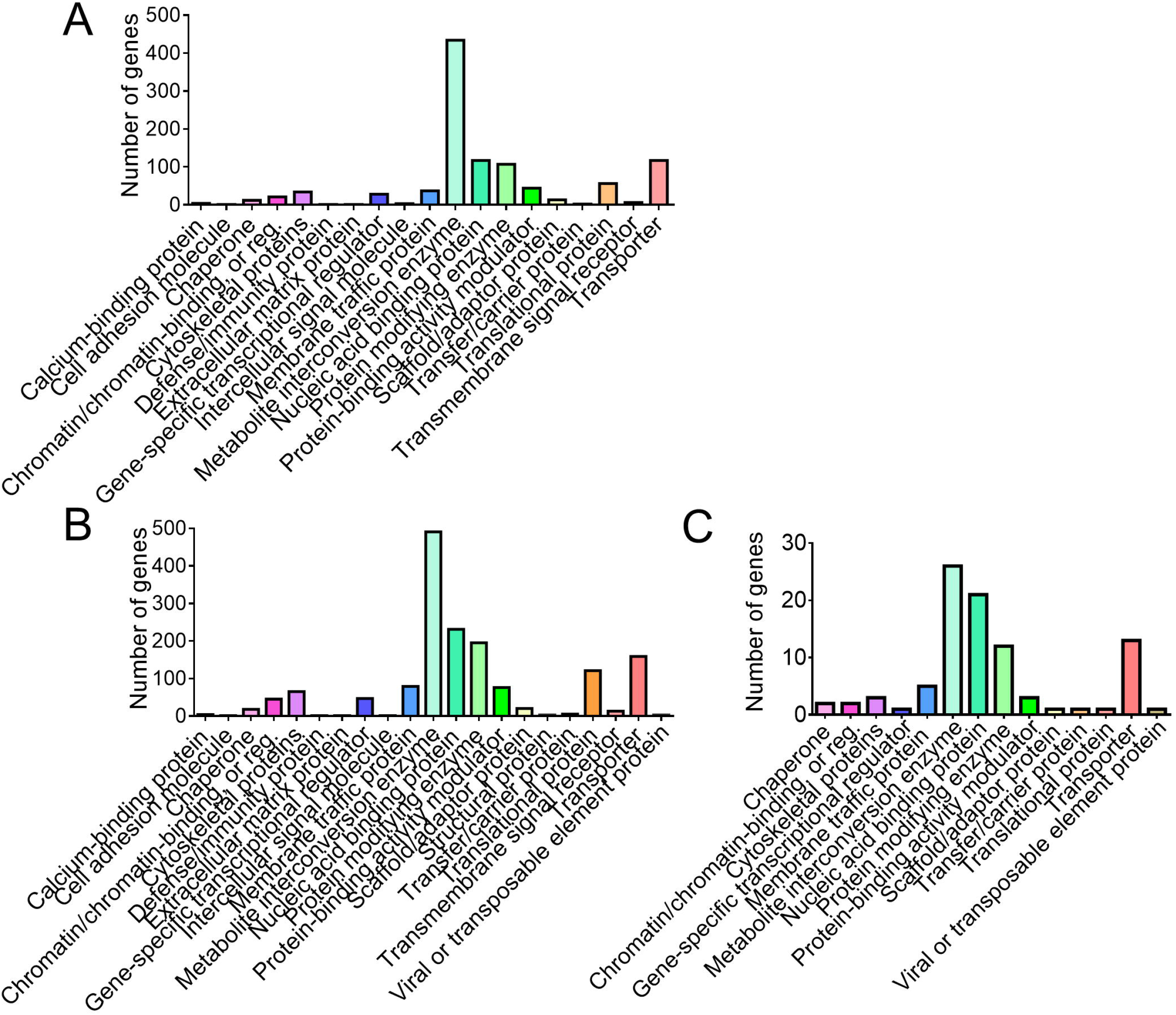
PQS can be found in genes encoding proteins involved in metabolite interconversion, protein modification, nucleic acid binding, and transporter activity. The number of genes containing PQS were quantified and protein classes associated with the genes were identified using PANTHER. The most highly represented groups are reported for **(A)** *A. fumigatus* Af293, **(B)** *C. neoformans* JEC21, **(C)** and *C. albicans* SC5314.

In all organisms, PQS could be found in the highest frequency in genes associated with metabolite interconversion enzymes. In *A. fumigatus*, the number of genes associated with metabolite interconversion enzymes was 3.7-fold higher than the next represented protein class (434 genes vs. 117 genes for nucleic acid binding proteins and transporters; Figure 9A). In *C. neoformans* the number of genes associated with these enzymes was 2.1-fold higher compared to nucleic acid binding proteins (491 genes vs. 231 genes, respectively; Figure 9B). In *C. albicans*, the difference in the number of PQS-containing genes associated with metabolite interconversion enzymes and nucleic acid binding proteins was much lower (26 genes vs. 21 genes, respectively; 1.2-fold; Figure 9C). Surprisingly, when categorising genes based on gene ontology terms, there was an almost identical distribution of genes involved in biological functions, molecular functions, and cellular components between species (Supplementary Material).

### PQS are found in genes linked to virulence, drug resistance, or key biological processes in *A. fumigatus, C. neoformans*, and *C. albicans*

G4s have been discovered in virulence genes from several pathogens and have arisen as a promising target for antimicrobial therapy and overcoming drug resistance [25]. Moreover, an iM in the promoter of the HIV-1 pro-viral genome has also been recently been described [31]. Thus, whether PQS could be found in genes associated with virulence/drug resistance in *A. fumigatus, C. neoformans*, and *C. albicans* was explored. Although the list is not exhaustive (there are many proteins still yet to be characterised), there were many interesting candidates that arose from the analysis. In total, PQS were found in over 100 genes associated with the virulence, drug resistance, or key biological processes of *A. fumigatus* (39 genes), *C. neoformans* (41 genes), and *C. albicans* (27; Tables 1-3).

**Table 1.**
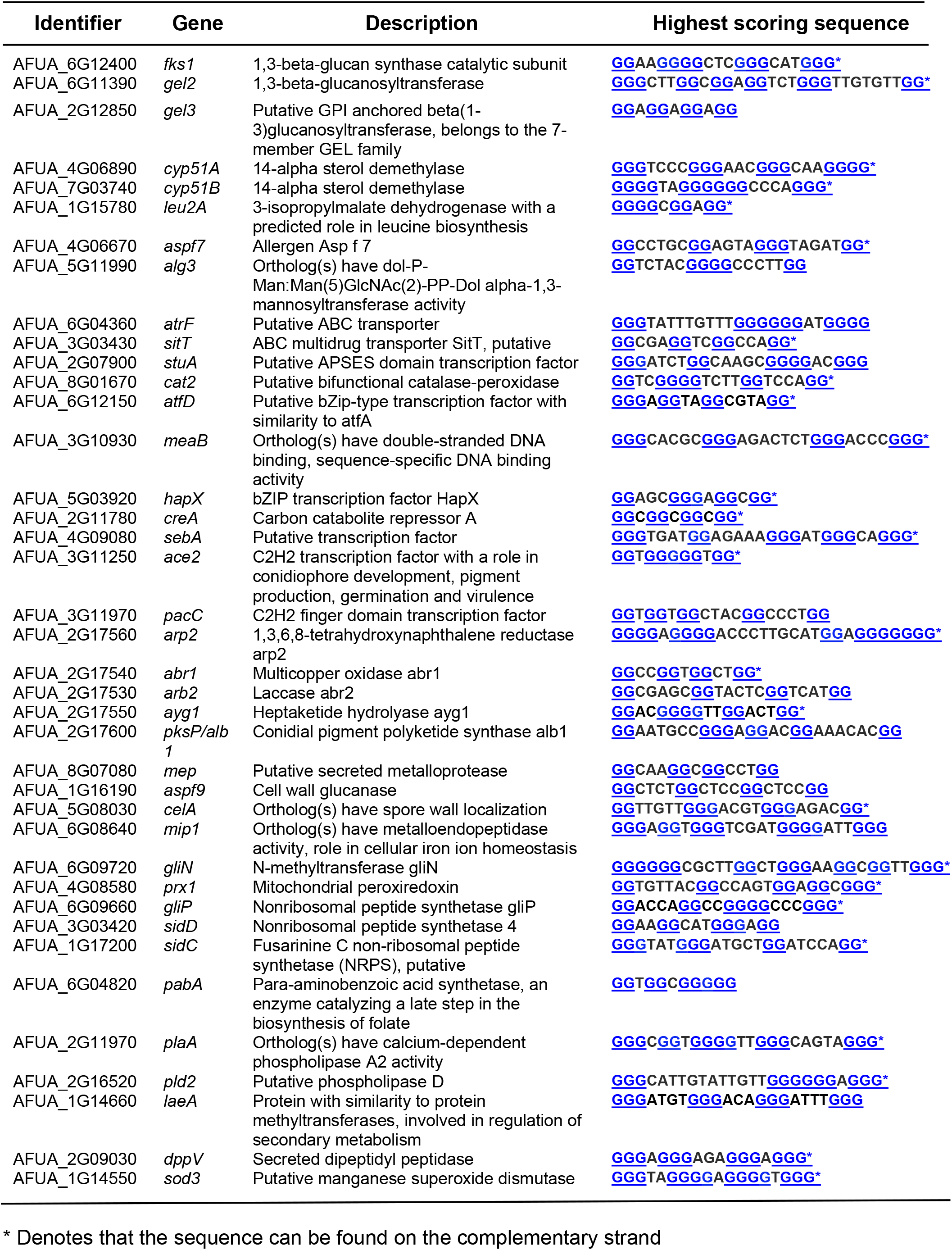
PQS containing genes in *A. fumigatus* linked to virulence, drug resistance, or key biological processes.

In *A. fumigatus*, PQS could be found in notable genes, including the 14-α sterol demethylases (*cyp51A* and *cyp51B*), the 1,3-β-glucan synthase catalytic subunit *fks1*, and ABC drug exporter *atrF*, which are involved in drug resistance. In addition to genes involved in virulence, including transcription factors *stuA, hapX*, and *pacC*, genes involved in pigment biosynthesis (*pksP, arp2, abr1, abr2*, and *ayg1*), a master regulator of secondary metabolism *laeA*, and *gliN and gliP* which are involved in the synthesis of gliotoxin (Table 1).

As PQS could be found in almost two-thirds of *C. neoformans* genes, it was not surprising that PQS could be found in those associated with virulence. These included the ABC transporter *afr1* (which is associated with fluconazole resistance), the protein kinases *fsk* and *hog1*, the calcineurin-associated genes *crz1* and *cna1, pacC/rim101* like in *A. fumigatus*, and numerous capsule-associated genes (the main virulence factor of *Cryptococcus*) including *cap2, cap5, cap10, cap59, cap60, cap64, cas31, cas33*, and *cxt1* (Table 2).

**Table 2.**
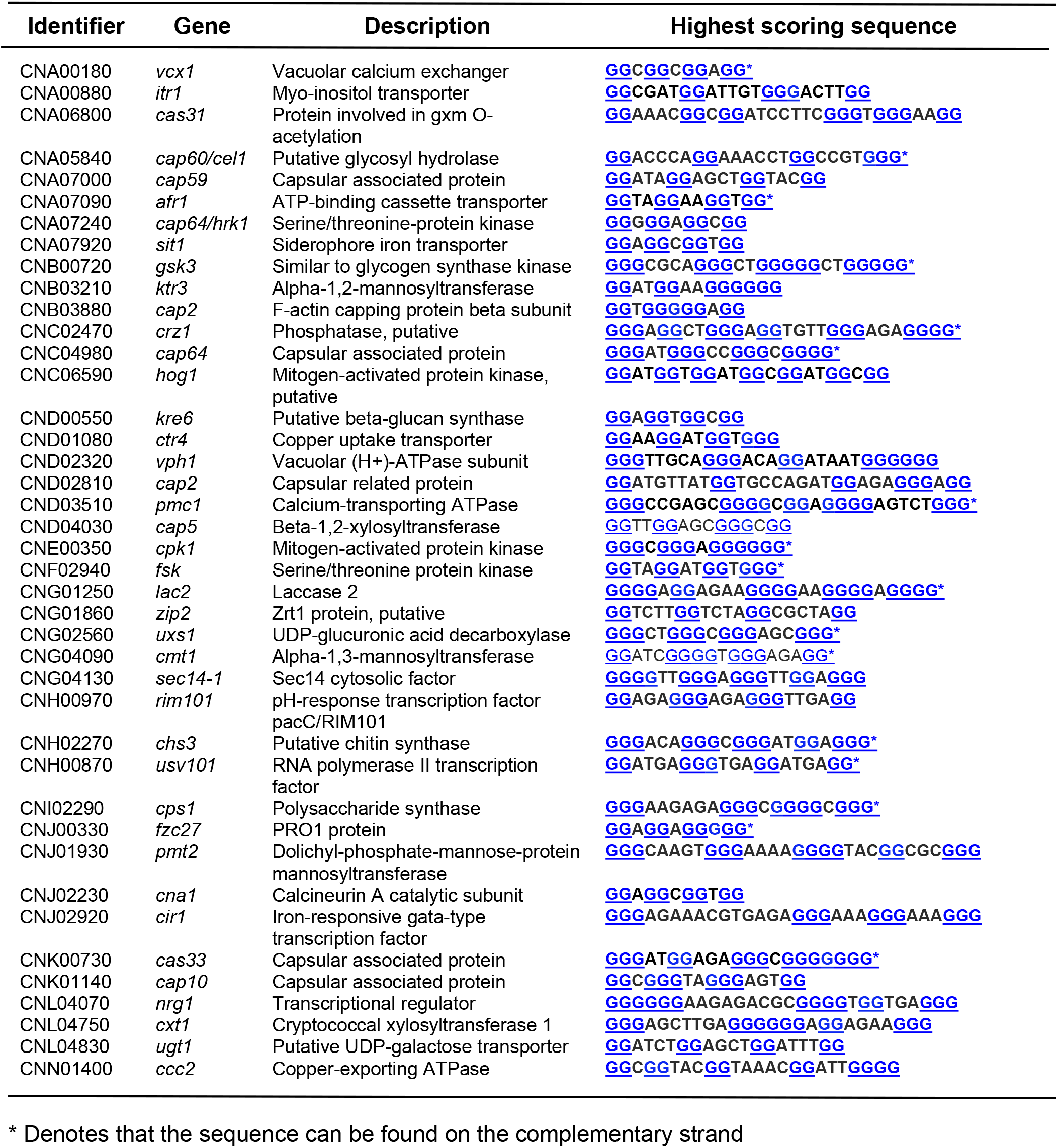
PQS containing genes in *C. neoformans* linked to virulence, drug resistance, or key biological processes.

There were very few genes in *C. albicans* that contained sequences likely to form quadruplexes, and thus, quadruplexes might be less important in this organism. Notable genes included the iron permeases *ftr1* and *ftr2*, and a gene associated with flucytosine resistance (*rrp9*; Table 3).

**Table 3.**
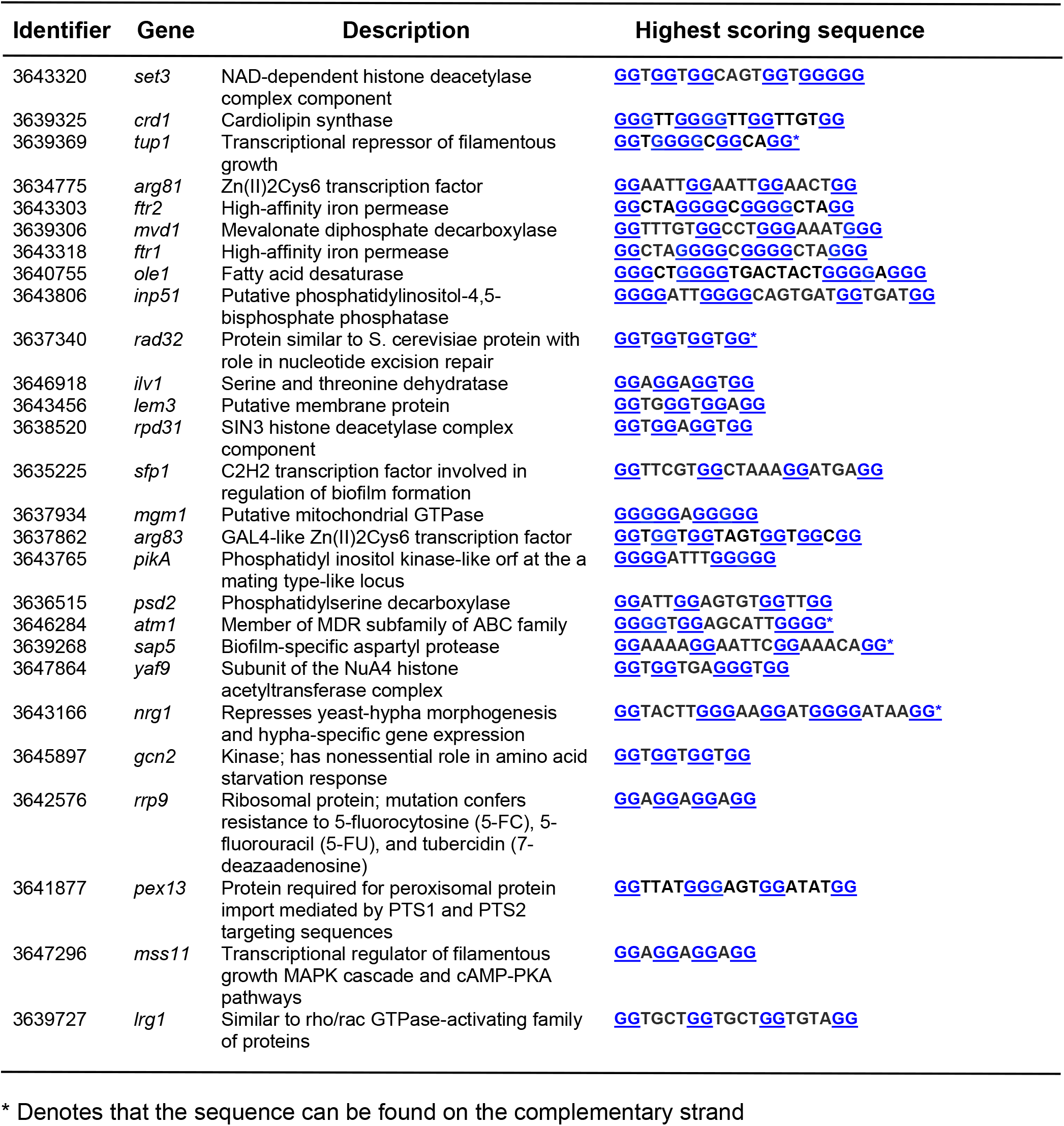
PQS containing genes in *C. albicans* linked to virulence, drug resistance, or key biological processes.

The highest scoring potential quadruplex-forming sequences for each of these genes were then re-analysed in an alternative PQS predictive algorithm called QGRS Mapper. In this instance, the scores of known quadruplex-forming sequences were compared to scores of the PQS in fungi. This was conducted to provide further insight into whether these sequences were likely to form quadruplex structures.

Known quadruplex-forming sequences (n=94) were shown to predominantly have QGRS scores of 21, 42, or 63 (depending on the number of G-tetrads – 2, 3, or 4, respectively; Supplementary Figures). The genes containing PQS from both *A. fumigatus* and *C. neoformans* had more sequences scoring between 21 and 42 compared to *C. albicans* and are thus more likely to form quadruplex structures (Supplementary Figures).

### Genes in *A. fumigatus* that are upregulated during fungal germination, in response to stress, and in biofilms are enriched in PQS compared to the whole genome

Although PQS could be found in important fungal genes, we aimed to link the presence of PQS within these organisms with potential biological and pathophysiological functions. To investigate this, transcriptome datasets of *A. fumigatus* during germination [39], hypoxia [40], iron limitation and oxidative stress [41], or in biofilms [43] were analysed. In each instance, the top 20 upregulated genes (compared to dormant/unstressed *A. fumigatus* controls) were investigated for the presence and frequency of PQS. This was achieved via examining the sequences in both QGRS Mapper and G4Hunter using the default search settings. Gene sequences which were identified as containing PQS by both algorithms were used for analysis.

Notably, PQS could be found in 77.5% of the genes investigated (Figure 10A). This included genes which were upregulated in germinating conidia (16/20; 80.0%), hyphae (14/20; 70.0%), after 12 hours of hypoxia (17/20; 85.0%), following iron limitation (17/20; 85.0%), undergoing oxidative stress (14/20; 70.0%), and in biofilms (15/20; 75.0%). These genes were investigated further to assess whether they were enriched with PQS and contained a higher frequency of PQS comparative to the average frequency found in the *A. fumigatus* genome. Using the default G4Hunter settings (window size 25 nt and threshold of 1.2) the PQS frequency of the *A. fumigatus* genome was 1.55 PQS/kbp (red dashed line; Figure 10B). In all cases, the average PQS frequencies in the upregulated genes were higher than the average PQS observed throughout the entire genome (Figure. 10B). The average PQS frequencies in upregulated PQS-containing genes were 2.97 PQS/kbp (germinating conidia), 3.72 PQS/kbp (hyphae), 2.80 PQS/kbp (12 h hypoxia), 2.66 PQS/kbp (iron starvation), 2.32 PQS/kbp (oxidative stress), and 2.26 PQS/kbp (biofilms; Figure 10B). Although, there were a range of PQS frequencies observed between the genes from 0.34 to 11.90 PQS/kbp. The genes containing the highest PQS frequencies for each condition were AFUA_8G01710 in germinating conidia and hyphae (11.90 PQS/kbp), AFUA_4G09580 in hypoxic fungi (5.59 PQS/kbp), AFUA_3G03650 during iron limitation (8.50 PQS/kbp), AFUA_5G10220 during oxidative stress (5.28 PQS/kbp), and AFUA_8G01980 in biofilms (5.90 PQS/kbp). Interestingly, each of these genes were upregulated in at least 3 out the 6 conditions investigated.

**Figure 10.**
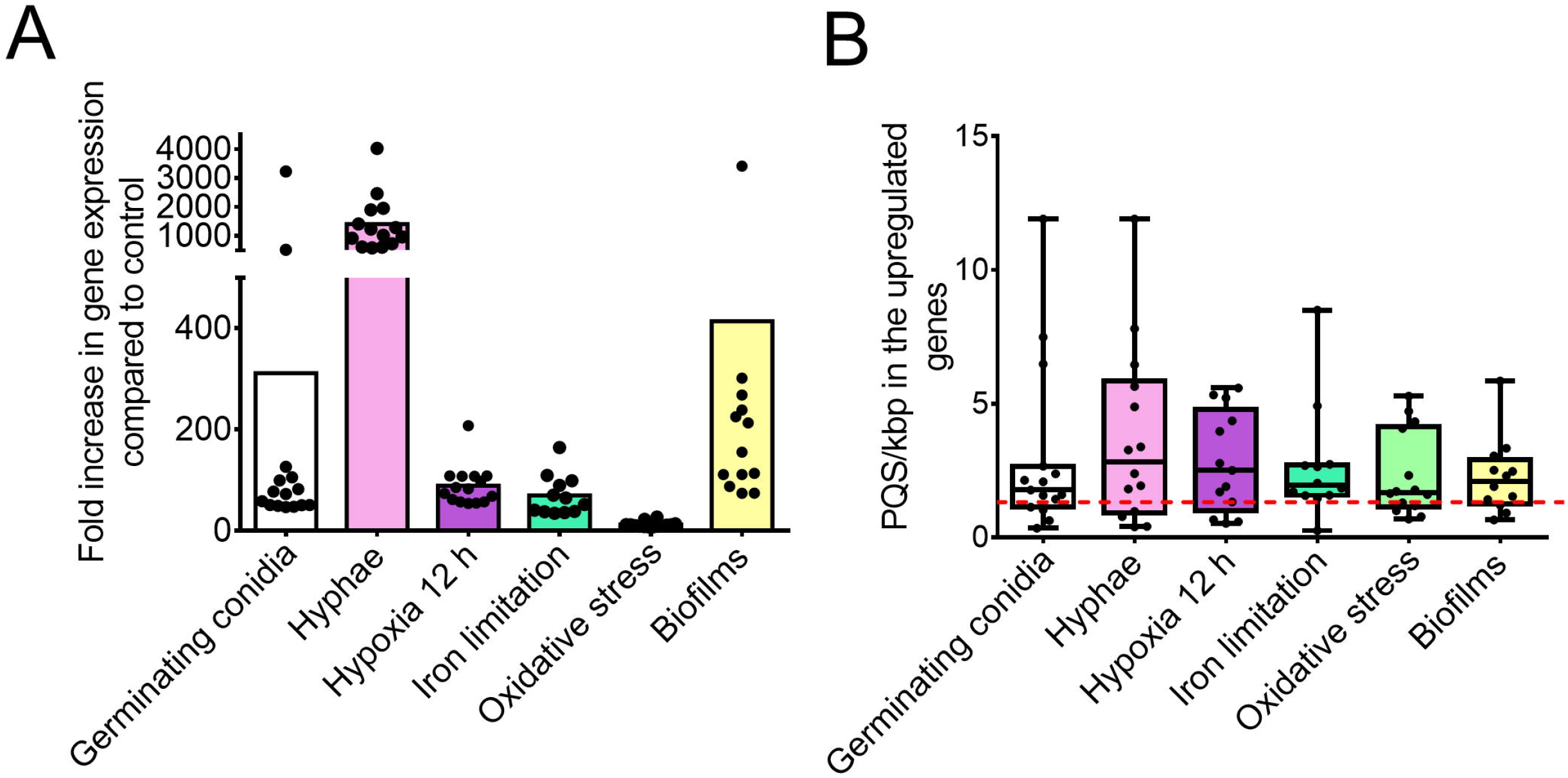
PQS can be found in genes which are highly upregulated during fungal germination, in response to environmental stresses, and in biofilms in *A. fumigatus*. Transcriptomes of *A. fumigatus* during germination, hypoxia, iron limiting conditions, oxidative stress, and in biofilms, were analysed and the 20 most upregulated genes were further investigated using QGRS Mapper and G4Hunter. (A) Genes that were highly upregulated (compared to dormant or untreated *A. fumigatus* conidia) contained PQS. Data points represent genes that contain PQS. (B) The PQS frequency in genes upregulated in these conditions is higher when compared to the average PQS/kbp for the entire genome (red-dashed line), analysed with the default G4Hunter settings (window size 25; threshold 1.2). Data points represent PQS frequency of an individual gene. Boxplot represents the median, maximum, and minimum values.

## Discussion

In this study, the number of potential quadruplex-forming sequences within the genomes of fungi were computationally predicted and their potential involvement in pathogenicity was discussed. Several important observations were made. This was the first study to identify the heterogeneity of PQS amongst genetically distinct fungal species. Moreover, we highlighted that pathogenic *Aspergillus* and *Cryptococcus* species contained fewer PQS compared to their non-pathogenic counterparts and these could be found throughout known genomic features, including genes, mRNA, repeat regions, tRNA, ncRNA, and rRNA. Genes containing PQS were associated with metabolism, nucleic acid-binding proteins, protein modifying enzymes, and transporters. Notably, PQS likely to form quadruplexes were identified in genes linked with fungal virulence or drug resistance, such as *cyp51A*, and could be found in genes upregulated during fungal growth and in response to stress.

The frequency of PQS throughout genomes is highly variable; for example, human genomes were shown to contain around 0.228 PQS/kbp, whereas the genomes of *Escherichia coli* contain around 0.028 PQS/kbp [15]. In this study we also found significant differences in the number and frequency of PQS throughout fungal genomes. For example, fungi from the Saccharomycotina (primarily composed of yeasts) contained a low frequency of PQS. Conversely, filamentous fungi from the Pezizomycotina (containing many important pathogens of both plants and humans) had much higher PQS frequencies and >15-fold more PQS than fungi from the Saccharomycotina on average. It has previously been shown that G4s contribute to genetic instability in yeasts (*Saccharomyces* spp.,) and as suggested for bacteria, it’s possible that G4s may also have been deselected through evolution in ascomycete yeasts [45, 46]. However, things are more complicated concerning basidiomycetous yeasts such as *Cryptococcus* spp. and *Malassezia* spp. Cryptococcal species such as *C. wingfieldii, C. amylolentus*, and *C. floricola* had some of the highest PQS frequencies in the study. Alternatively, all the *Malassezia* species investigated had much lower PQS frequencies than expected, which was surprising given their genomes were GC rich compared to many other fungi (>55% GC content).

Notably, the genomes of pathogenic aspergilli and cryptococci contained significantly lower PQS frequencies relative to their non-pathogenic counterparts. This trend was observed throughout the Pezizomycotina overall. Conversely, there was no correlation between PQS and pathogenicity between *Candida* species or within the Saccharomycotina. The exact reasons underlying the loss of PQS are not currently known. Fungi may have evolved to retain the most essential quadruplex-forming sequences or those that provided them with a pathogenic advantage. In the case of *Aspergillus* and *Cryptococcus* species, this loss of PQS might be linked with evasion of the immune response or enhanced pathogenicity. Furthermore, the observed loss of PQS within *Candida* and *Malassezia* species might be associated with their ability to form commensal relationships and avoid stimulating a host response. Interestingly, loss of PQS has recently also been observed in pathogenic *Coronaviridae* [47]. It has also been reported that host nucleolin (an RNA-binding protein) can bind and stabilise quadruplexes in the LTR promoter of HIV-1, which can silence viral transcription [48]. Therefore, in this situation, loss of quadruplexes would be beneficial for immune evasion. Contrarily, it has been suggested that iMs within HIV-1 are triggered after they promote the acidification of intracellular compartments, modulating viral processes [31]. In this instance, the formation of quadruplexes could be beneficial and promote pathogenicity. The importance of quadruplex stabilisation may therefore be situational and could be detrimental or beneficial depending on the environment. It is known that fungal pathogens such as *A. fumigatus* and *C. neoformans* can survive the acidic conditions within intracellular organelles to propagate infection [49, 50]. Therefore, it could be interesting to investigate the dynamics/flux of quadruplex formation under conditions that fungi would experience within the host and whether they contribute to fungal survival and propagation.

Here, we identified many PQS within the CDS, genes, and mRNA of *A. fumigatus, C. neoformans*, and *C. albicans*; but a higher frequency of PQS within the repeat regions of *A. fumigatus*, and the repeat regions and rRNA of *C. albicans*. It has been observed that *S. cerevisiae* promoters and open reading frames are enriched with sequences capable of potentially forming intramolecular G4s [51]. A recent study by Čutová *et al*., also demonstrated that inverted repeats were enriched in the centromeres and rDNA/rRNA regions, and G4s were enriched in the telomeres and tRNAs of *S. cerevisiae* [52].

There were large differences between the numbers of genes containing PQS in these organisms, but PQS were found distributed amongst genes encoding proteins from similar classes. The highest represented groups being metabolite interconversion enzymes, nucleic acid binding proteins, transporters, and protein modifying enzymes in all. Furthermore, genes containing PQS within *S. cerevisiae* promoter regions were primarily involved in metabolism. Which supports our observations and is equally concordant with observations made in both humans and bacteria [53, 54]. It is widely acknowledged that metabolism in fungi is central to its virulence, pathogenicity, and survival [55]. The ability of a fungus to rapidly adapt to the host microenvironment can be achieved through such processes as metabolic remodelling, stress resistance, and the utilisation of amino acids [56]. This in turn has a significant effect on the triggering of important virulence traits, like the production of hyphae, biofilm growth, capsule formation in *C. neoformans*, and melanisation [56]. Moreover, metabolic pathways influence fungal vulnerability to innate immune defences and can regulate susceptibility to antifungal drugs [57]. Not only primary metabolism, secondary metabolism results in the release of various secondary metabolites and toxins, such as gliotoxin, aflatoxin, and candidalysin [58, 59]. Interestingly, PQS could also be found in many of the most upregulated genes during germination, in response to environmental stress, or in biofilms. Additionally, these upregulated genes were enriched in PQS. Thus, the enrichment of PQS within these genes is suggestive of the potential importance of quadruplexes in the regulation of biological functions and pathogenicity in fungi, and it would be interesting to explore the effects quadruplex-targeting ligands could have on these pathways.

PQS could also be found within key genes linked to virulence and drug resistance in these fungi. In *A. fumigatus*, PQS could be found in *cyp51A, cyp51B, atrF*, and *fsk1* (involved in resistance to azoles and echinocandins) [60-62], *laeA, gliN and gliP*, (involved in toxin and metabolite biosynthesis) [63, 64], *pksP, arp2, abr1, arb2*, and *ayg1* (involved in the production of the virulence factor DHN-melanin) [65], in addition to numerous important transcription factors (*meaB, hapX, pacC*) [66]. Similarly, in *C. neoformans*, PQS could be found in genes linked to azole resistance (*afr1*) [67], melanin production (*lac2*) [68], and transcription factors (*pacC/rim101)*. The most notable virulence factor of *C. neoformans* is its polysaccharide capsule and PQS could be found in numerous capsule-associated genes (*cas31, cap60, cap59, cap64, cap2, cap5, cas33, cap10*, and *cxt1*) [69]. In *C. albicans* PQS could be found in genes such as the iron permeases *ftr1* and *ftr2* [70]. Notably, many of these genes contained PQS which have previously been shown to be capable of forming *bona fide* quadruplexes, such as the sequence <u>GG</u>A<u>GG</u>A<u>GG</u>A<u>GG</u> [71]. It is also interesting to highlight that these organisms contained many more G_2+_L_1-12_ compared to G_3+_L_1-12_ PQS sequences, which is a characteristic shared with *S. cerevisiae* [15].

There are now an ever-increasing number of G4s identified within genes linked to microbial pathogenicity. G4-forming motifs located in the *hsdS, recD*, and *pmrA* genes of *S. pneumoniae*, and *var* genes of *P. falciparum* have been identified to modulate host-pathogen interactions [72, 73]. Moreover, targeting G4s in *espK, espB*, and *cyp51* from *Mycobacterium tuberculosis* with the G4/iM-binding ligand TMPyP4 could inhibit transcription of these essential virulence genes [74]. Binding of several G4-targeting ligands to G4s in viruses have been shown to limit the virulence of the Herpes simplex virus-1, HIV-1, Ebola, and Hepatitis C [75]. Interestingly, dinuclear ruthenium (II) complexes (well-characterised G4 and iM DNA binding agents) have been shown to be active against methicillin-resistant *S. aureus* and vancomycin-resistant *Enterococcus* spp [29, 30]. Notably, another G4/iM-ligand, berberine, has also demonstrated activity towards fluconazole-resistant *C. albicans* and *C. neoformans* [76]. Thus, quadruplexes may provide a novel target to overcome the issues surrounding drug resistance.

## Conclusion

Identifying PQS within key virulence/drug resistance genes of fungal pathogens has highlighted structures that may be targeted by G4/iM-binding drugs to ameliorate fungal pathogenicity. However, we first need to identify whether these structures are formed and if they can modulate biological functions. Understanding whether quadruplexes could form under pathophysiological conditions, whether fungi themselves could release quadruplex-binding ligands to modulate host responses, or identifying whether there are fungal specific quadruplex-forming sequences could also be important factors to consider. Thus, targeting quadruplexes in fungi could present a novel target for the amelioration of fungal infection and drug resistance, although a more thorough investigation is necessary.

## Acknowledgements

V. B was supported by the Czech Science Foundation (18-15548S). I would like to acknowledge Phil Spence for contributing the G4 and iM structures and for critical reviewing of the manuscript. S.B conceived the study. E. F. W., N. B., V. B., and S.B., performed the data analysis. E. F. W. and S. B wrote the manuscript in consultation with Z. A. E. W. All authors contributed to interpretation of the results, provided critical feedback, and helped to shape the research and manuscript.

## Supplementary Figure Legends

**Supplementary Figure 1. (A-G)** A breakdown of PQS frequencies per kilobase pair (kbp) for fungal genera from the Ascomycota. Each individual dot represents a species within the genera.

**Supplementary Figure 2. (A-G)** A breakdown of PQS frequencies per kilobase pair (kbp) for fungal genera from the Basidiomycota. Each individual dot represents a species within the genera.

**Supplementary Figure 3.** A breakdown of PQS frequencies per kilobase pair (kbp) for fungal genera from the **(A)** Mucoromycota, **(B)** Zoopagomycota, **(C)** Chytridiomycota, **(D)** Microsporidia, and **(E)** Cryptomycota. Each individual dot represents a species within the genera.

**Supplementary Figure 4.** The frequency of PQS per kilobase pair (kbp) **(A)** and relative to GC content **(B)** in *Aspergillus* species. Blue bars represent non-pathogenic species, whilst pink bars represent pathogenic species.

**Supplementary Figure 5.** The frequency of PQS per kilobase pair (kbp) **(A)** and relative to GC content **(B)** in fungi from the Pezizomycotina that are pathogenic in animals or plants, or non-pathogenic. Points represent individual species. Error bars represent the SD. * indicates a significant difference (p<0.05).

**Supplementary Figure 6.** The frequency of PQS per kilobase pair (kbp) **(A)** and relative to GC content **(B)** in *Cryptococcus* species. Blue bars represent non-pathogenic species, whilst pink bars represent pathogenic species.

**Supplementary Figure 7.** The frequency of PQS per kilobase pair (kbp) **(A)** and relative to GC content **(B)** in *Candida* species. Blue bars represent non-pathogenic species, whilst pink bars represent pathogenic species.

**Supplementary Figure 8.** The frequency of PQS per kilobase pair (kbp) **(A)** and relative to GC content **(B)** in fungi from the Saccharomycotina that are pathogenic in animals or plants, or non-pathogenic. Points represent individual species. Error bars represent the SD. ns indicates no significant difference (p>0.05).

**Supplementary Figure 9.** The distribution and number of genes involved in biological processes in **(A and D)** *A. fumigatus*, **(B and E)** *C. neoformans*, and **(C and F)** *C. albicans*.

**Supplementary Figure 10.** The distribution and number of genes involved in molecular functions in **(A and D)** *A. fumigatus*, **(B and E)** *C. neoformans*, and **(C and F)** *C. albicans*.

**Supplementary Figure 11.** The distribution and number of genes involved in cellular components in **(A and D)** *A. fumigatus*, **(B and E)** *C. neoformans*, and **(C and F)** *C. albicans*.

**Supplementary Figure 12. The scores of PQS found in fungi compared to known quadruplex forming sequences.** PQS in the identified genes were scored in QGRS Mapper and compared against the scores of known quadruplexes (n=94) to predict the propensity of PQS sequences to form quadruplex structures. Sequences with scores of 21, 42, and 63 in QGRS Mapper could form G4s containing 2, 3, and 4-tetrads, respectively. Sequences containing G_2_+L_1-12_ generally produced scores of 20 or 21, while those containing G3+L1-12 produced scores between 38-42. F and R refer to the coding and complementary strands, respectively.

