## Supplementary figures and images for "Cross kingdom analysis of putative quadruplex-forming sequences in fungal genomes: novel antifungal targets to ameliorate fungal pathogenicity?"

### Supplementary Figure 1A-C

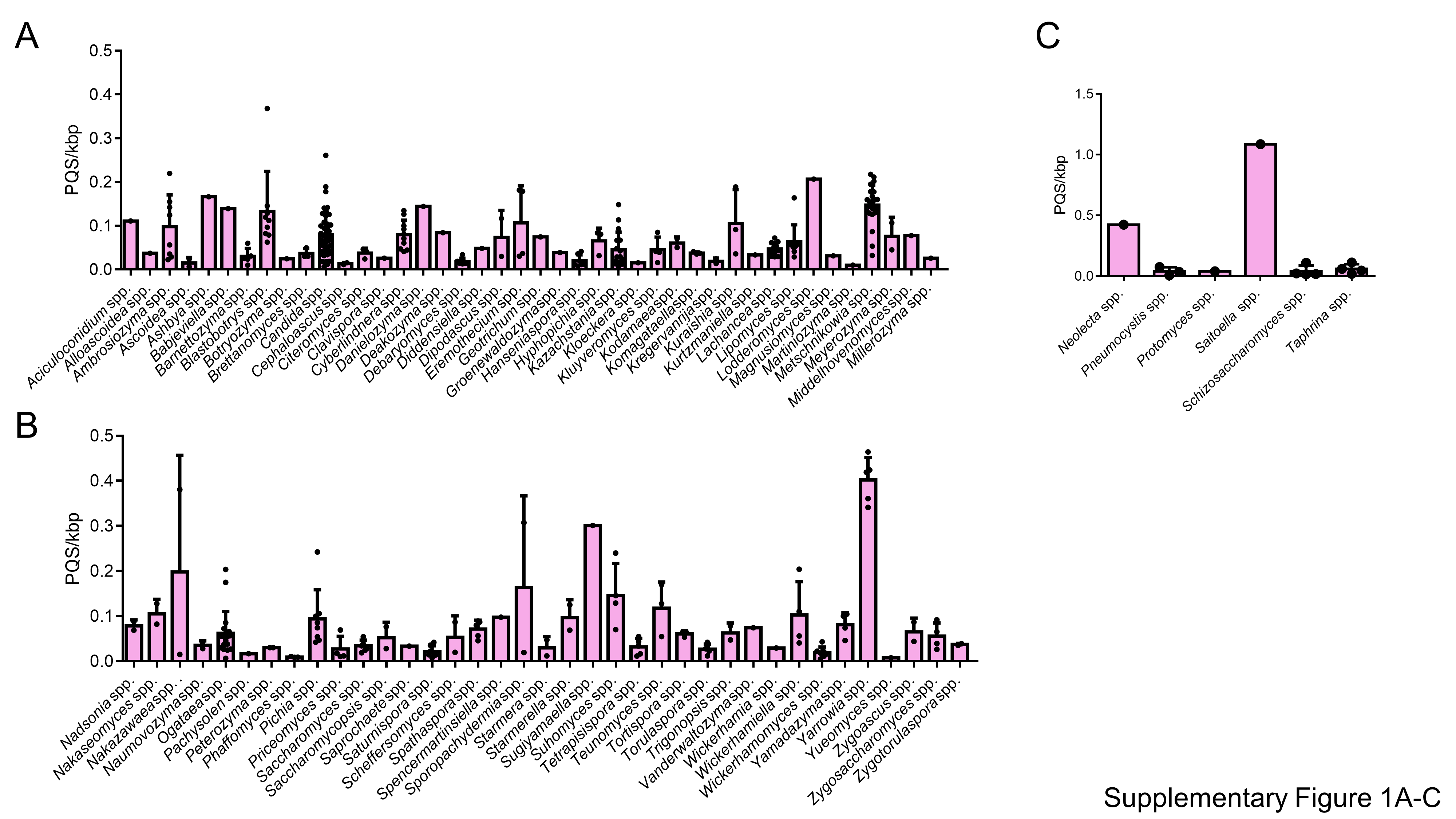

### Supplementary Figure 1D and E

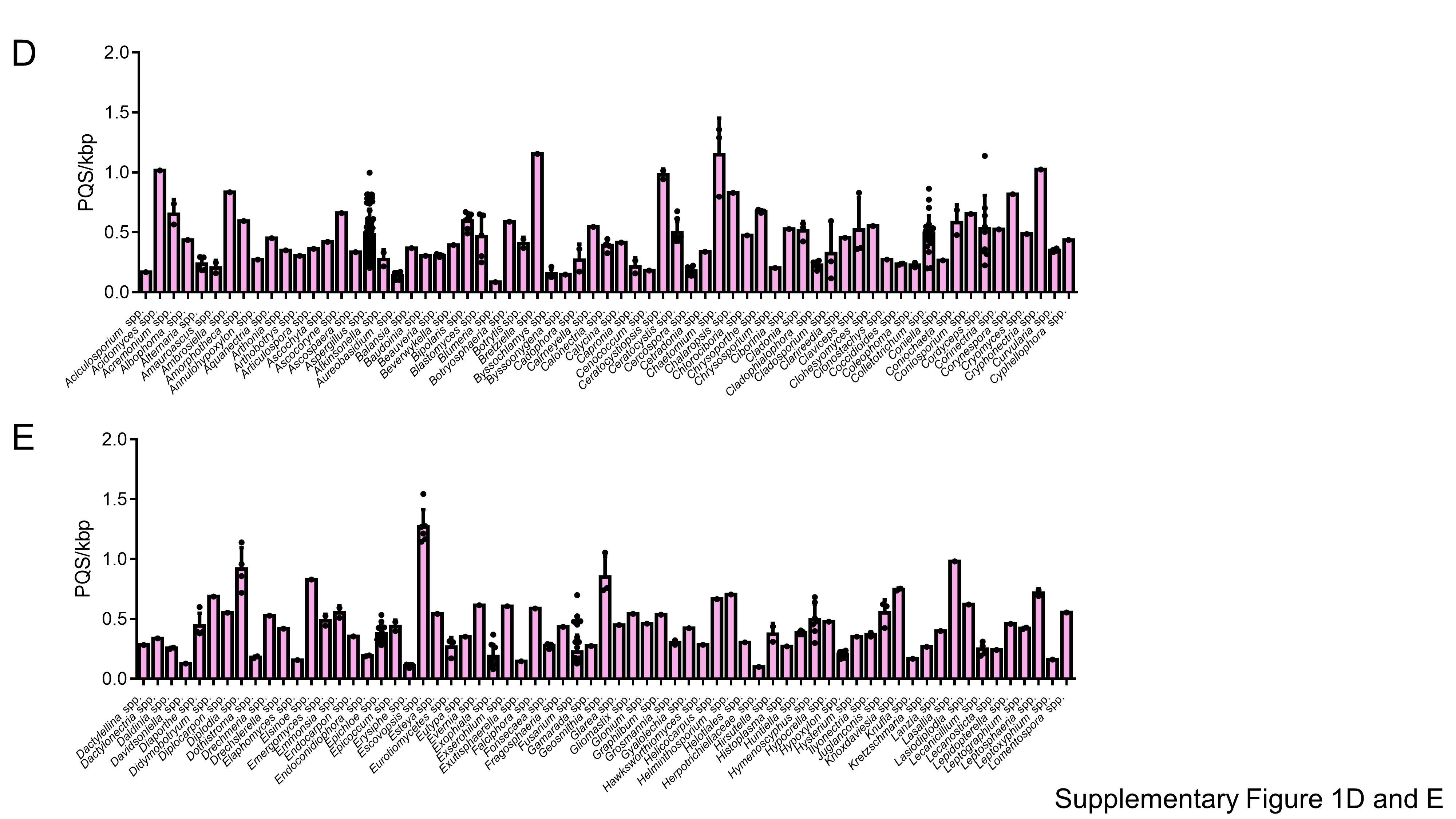

### Supplementary Figure 1F and G

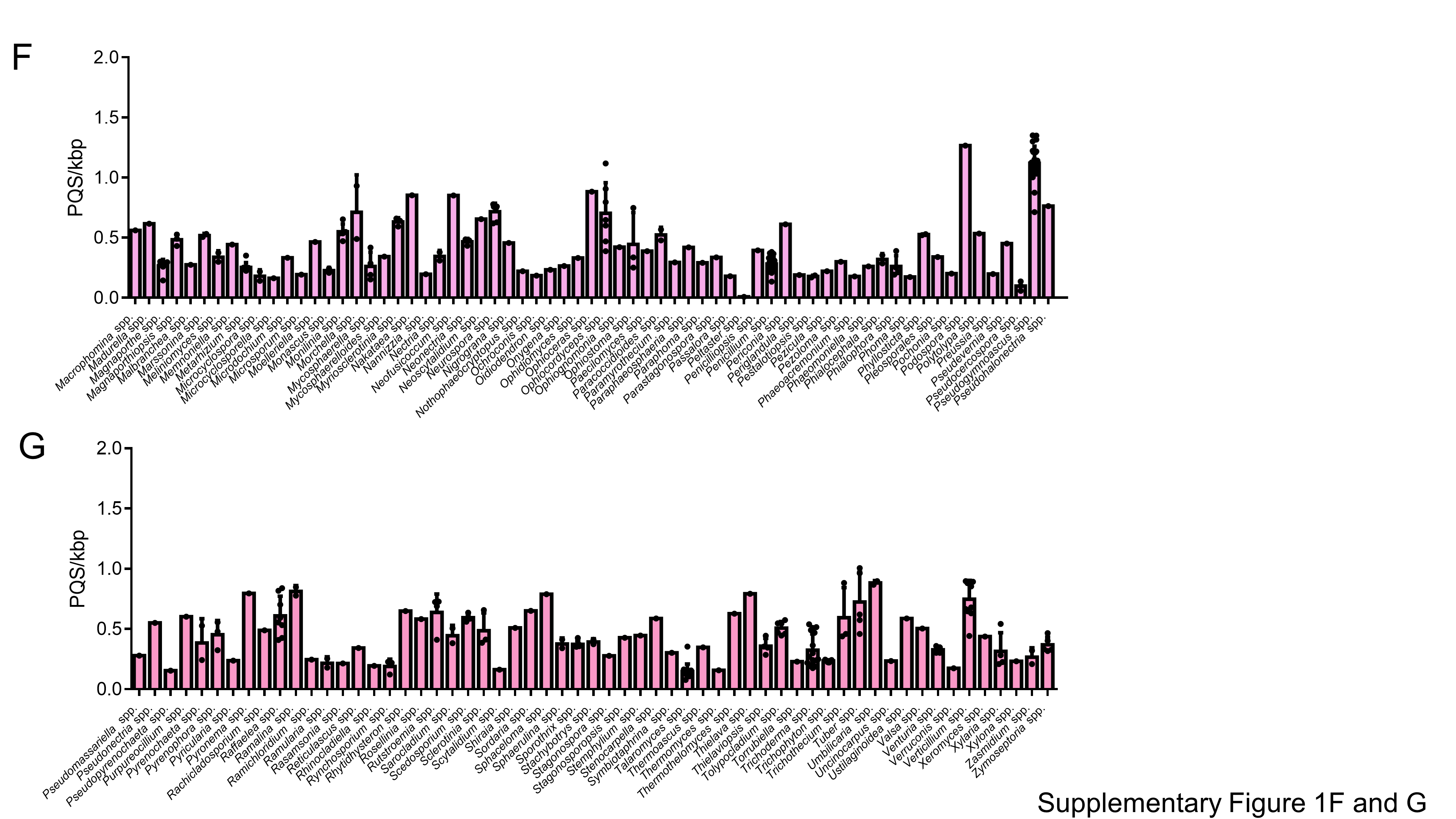

### Supplementary Figure 2

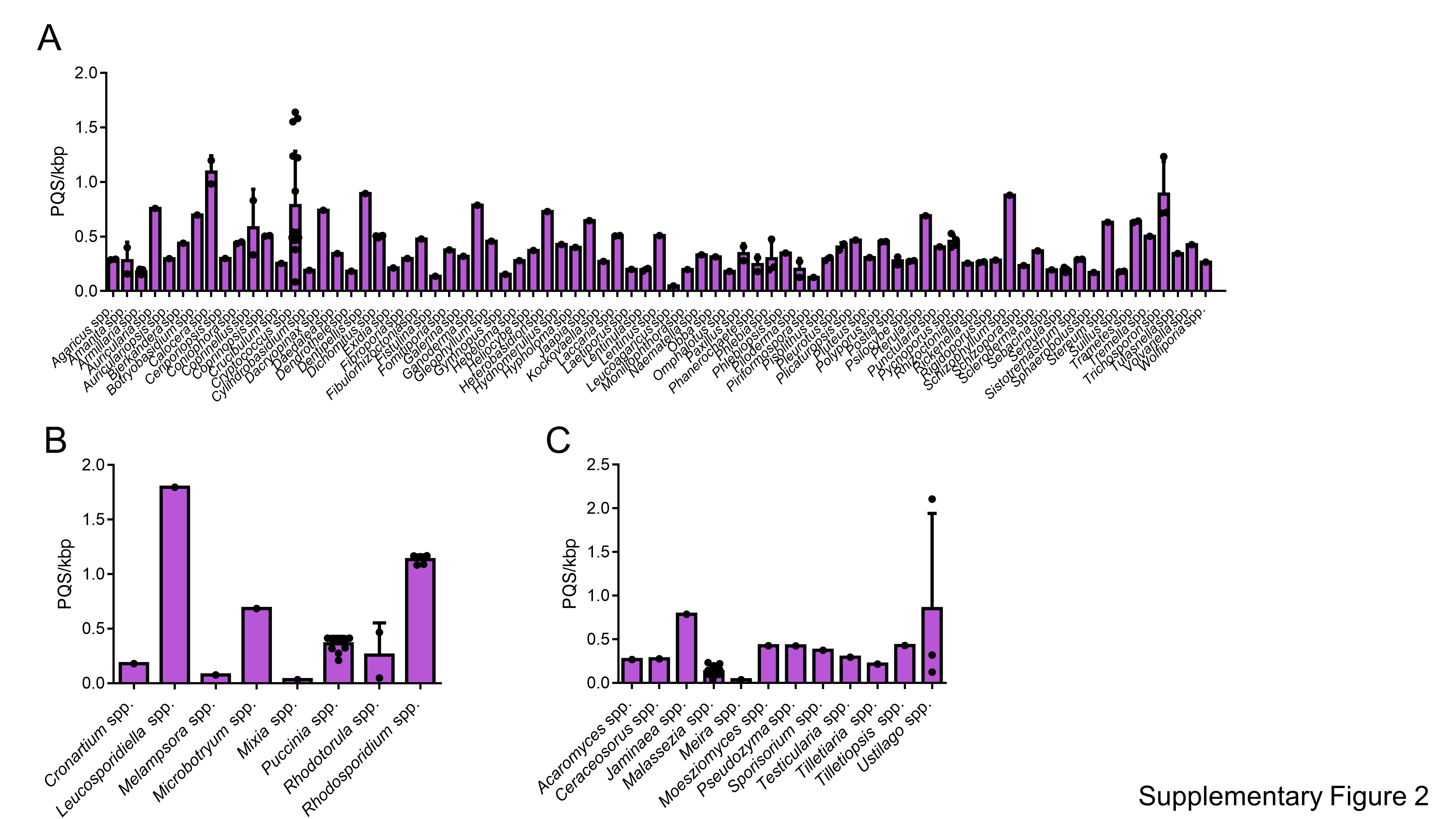

### Supplementary Figure 3

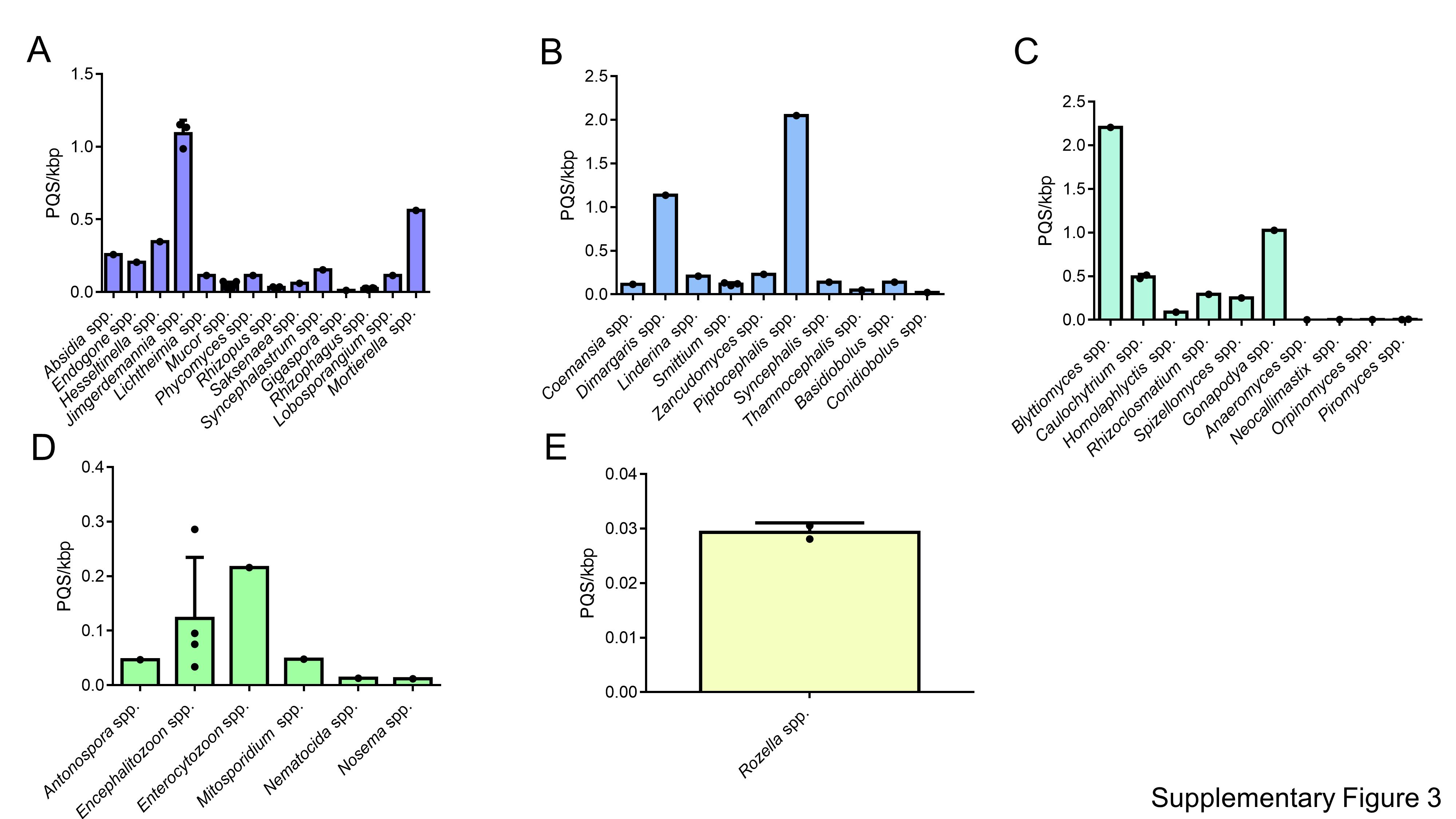

### Supplementary Figure 4

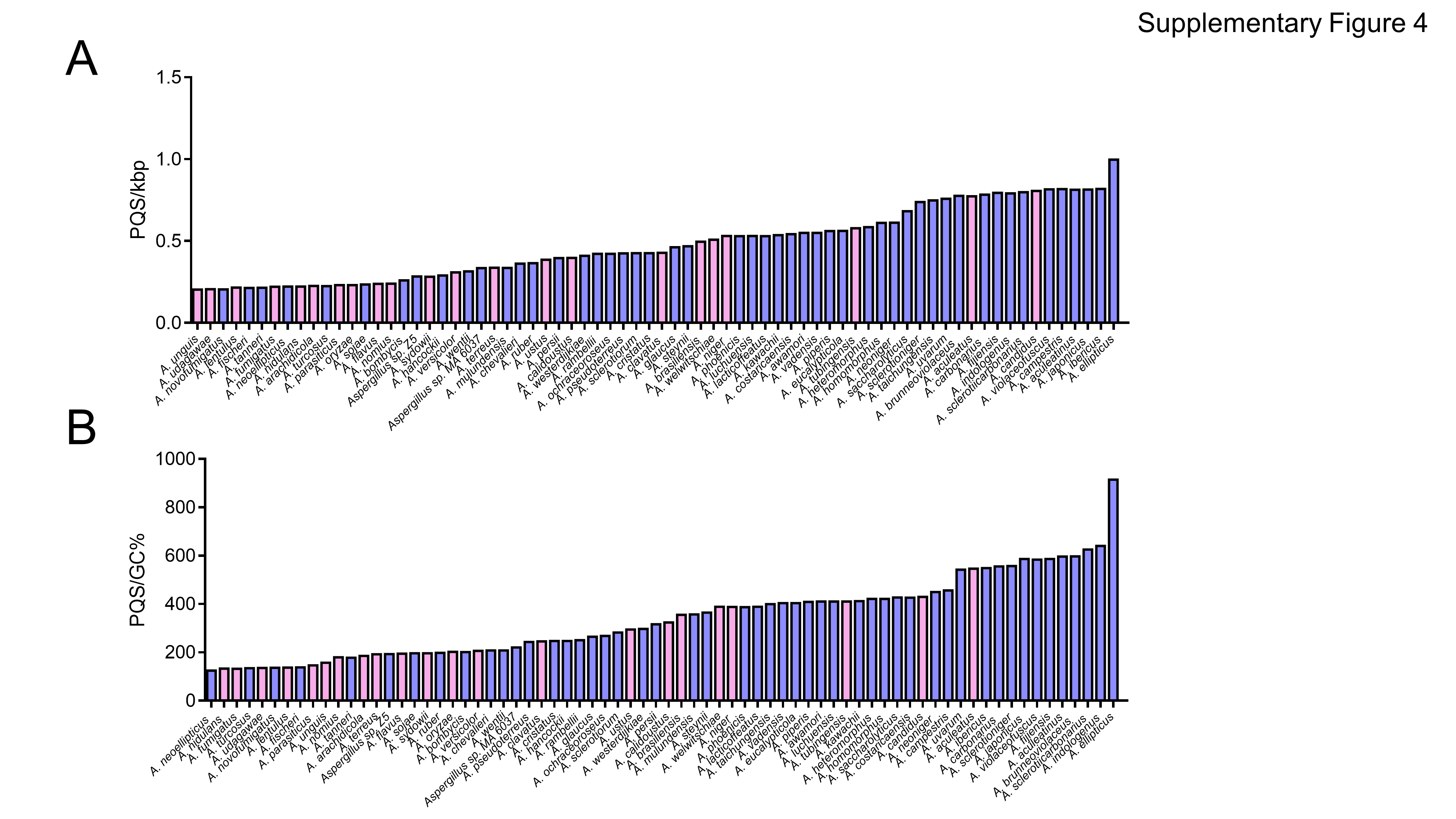

### Supplementary Figure 5

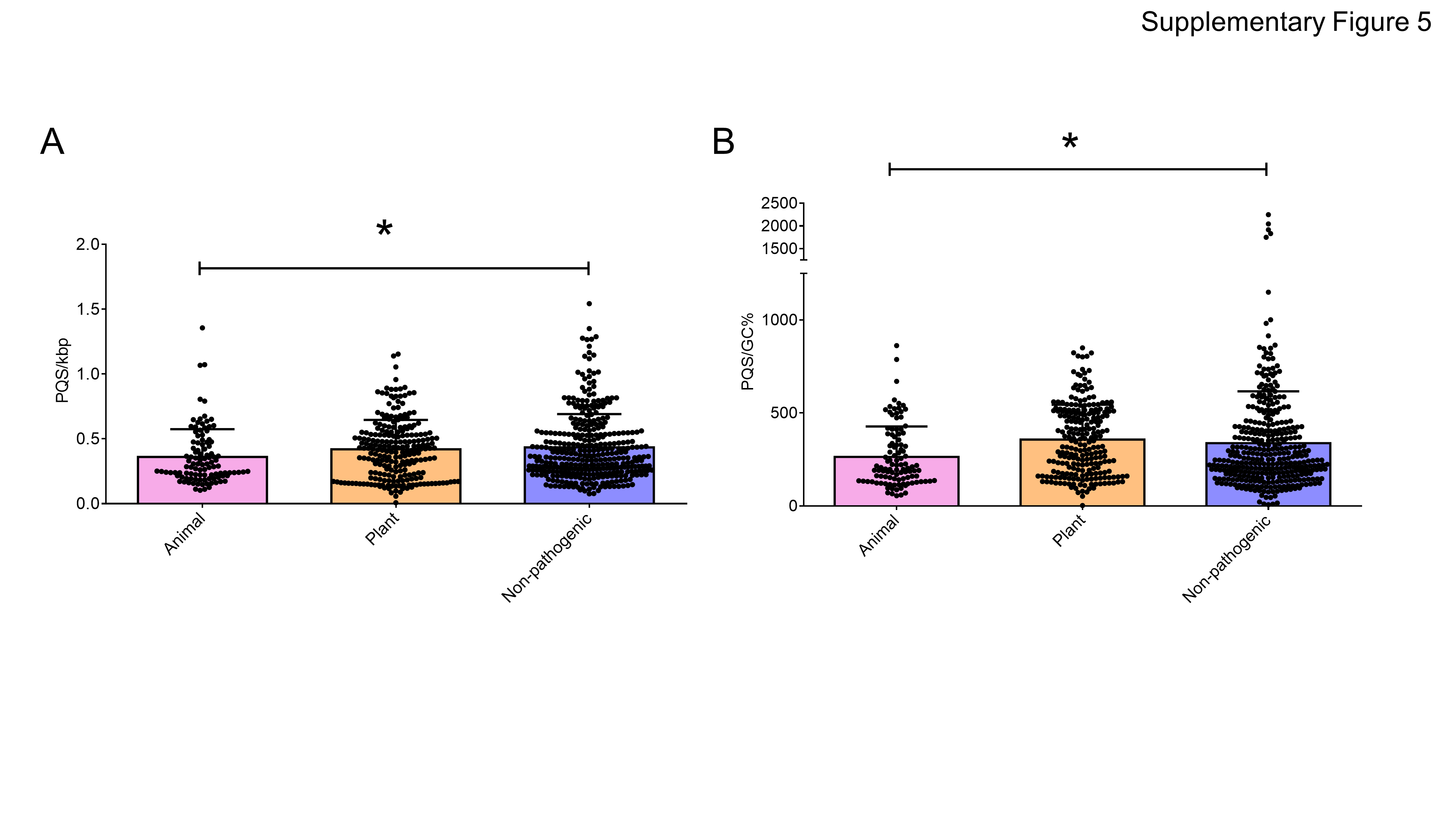

### Supplementary Figure 6

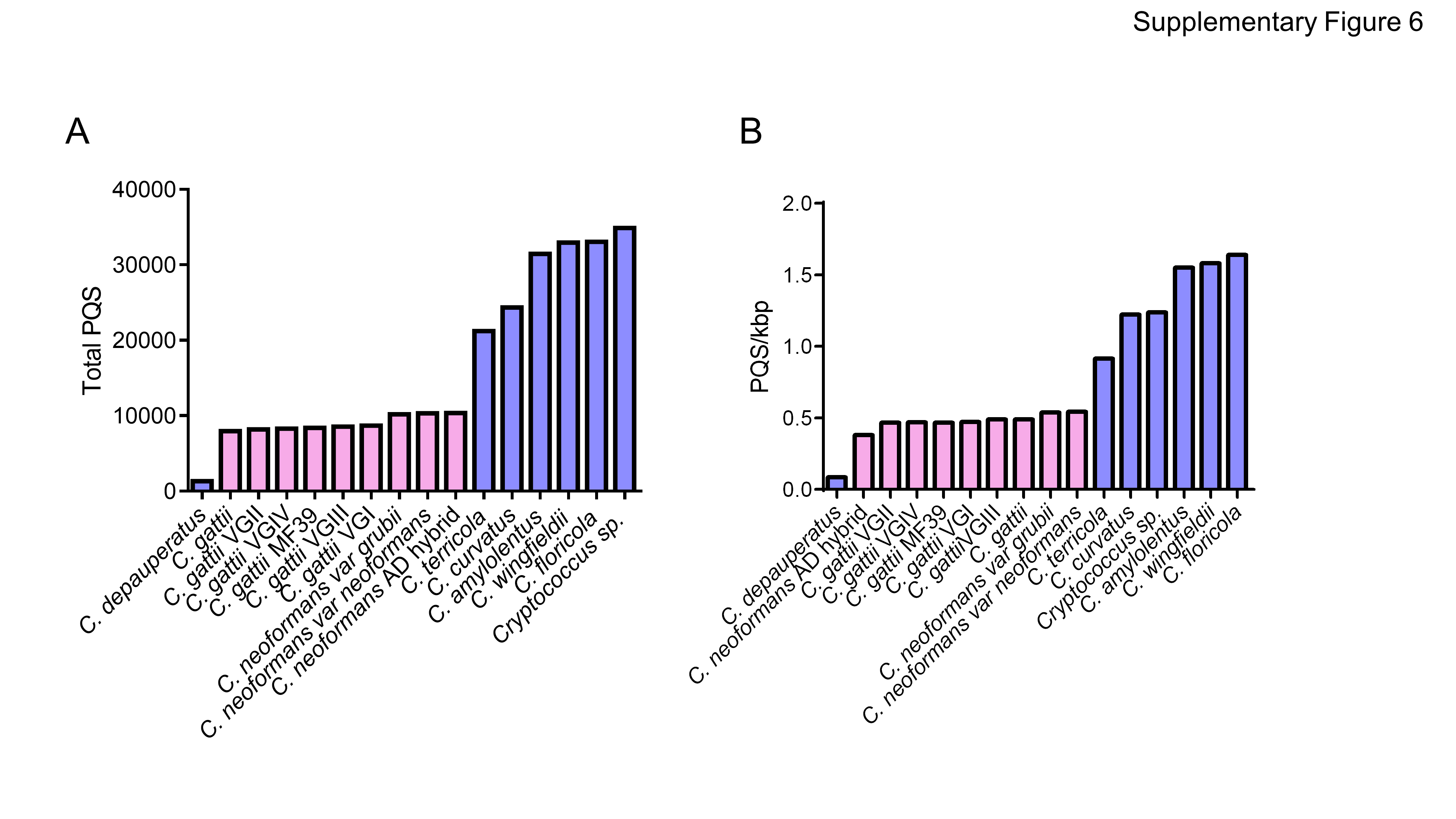

### Supplementary Figure 7

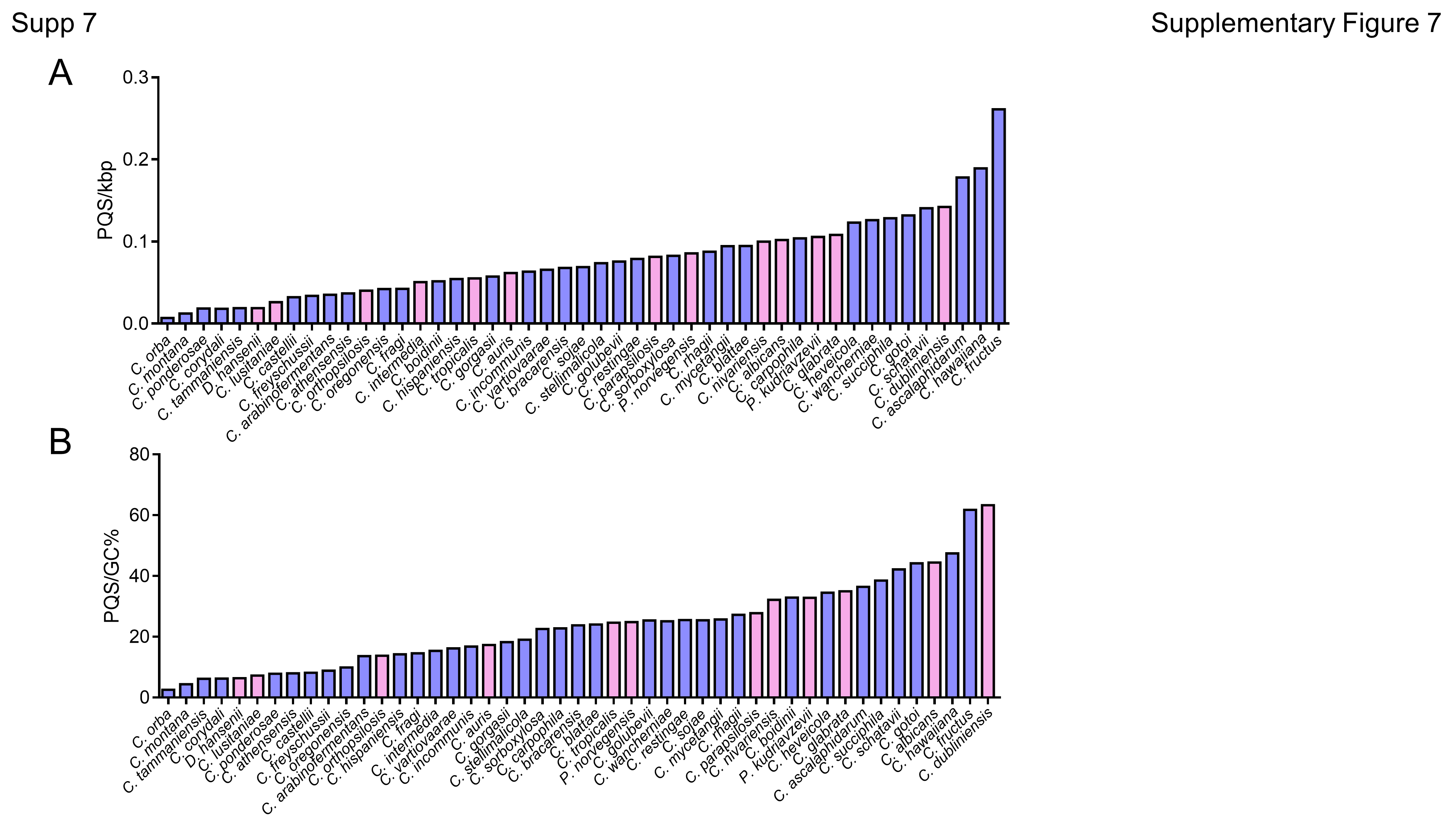

### Supplementary Figure 8

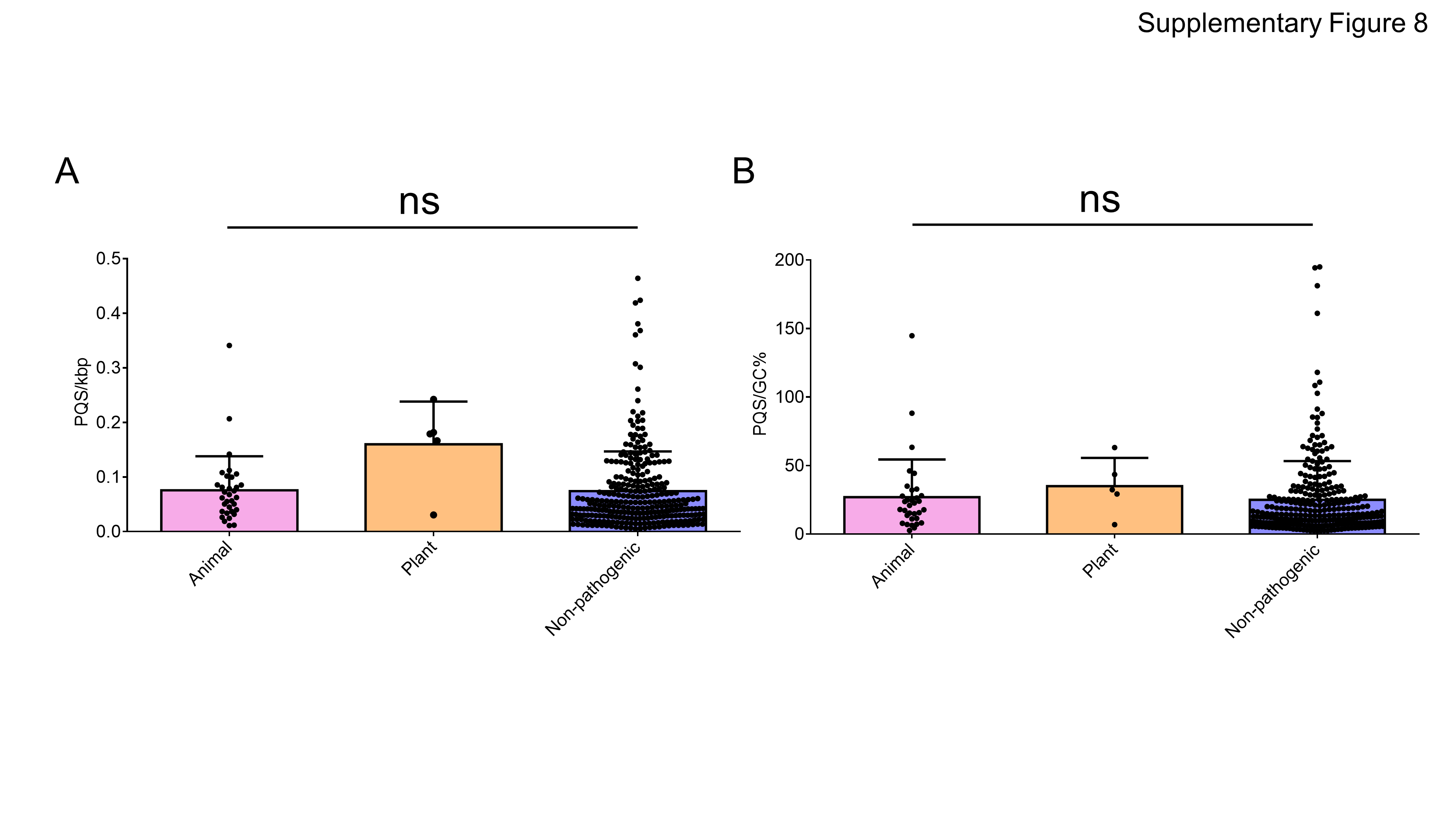

### Supplementary Figure 9

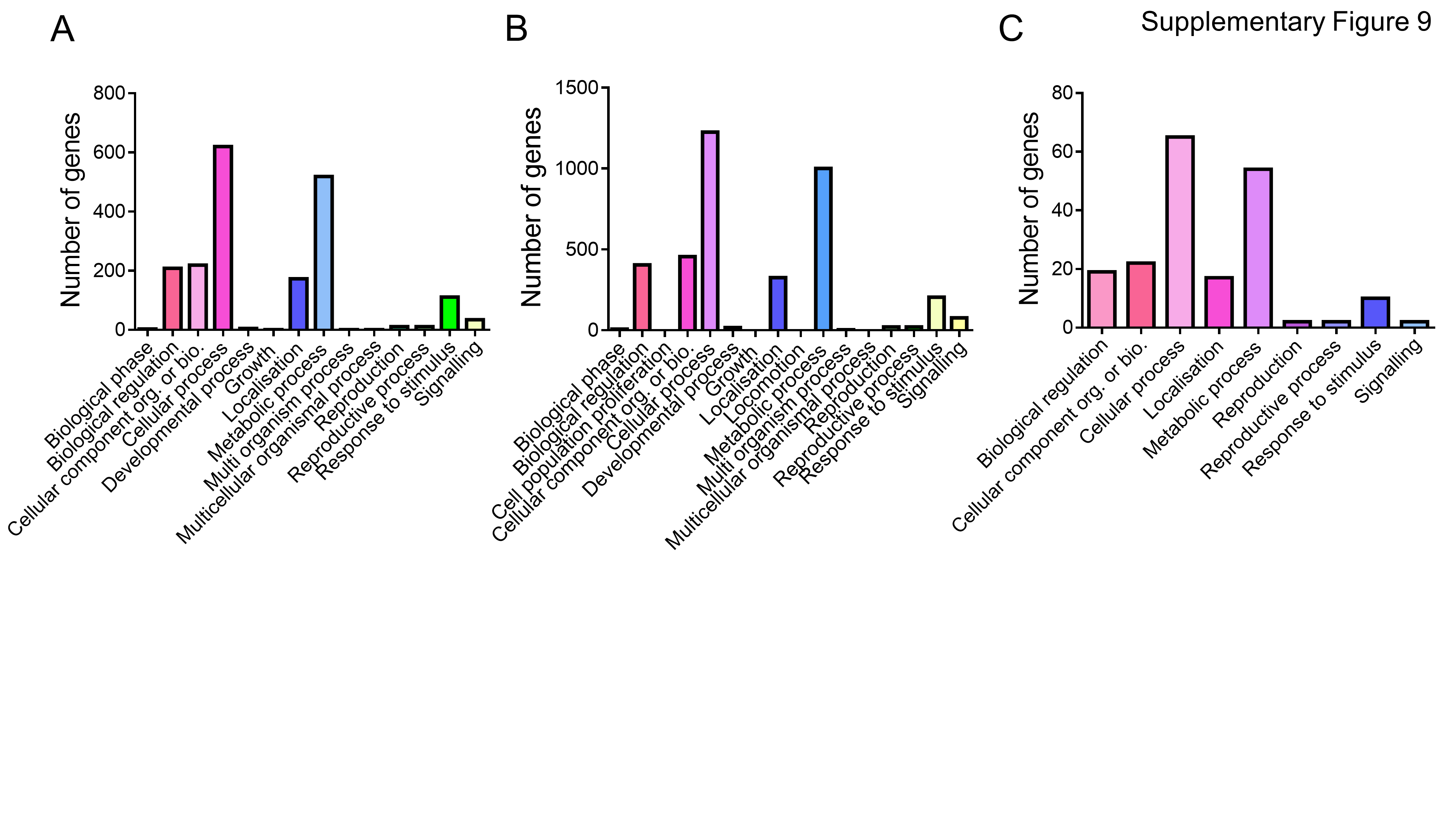

### Supplementary Figure 10

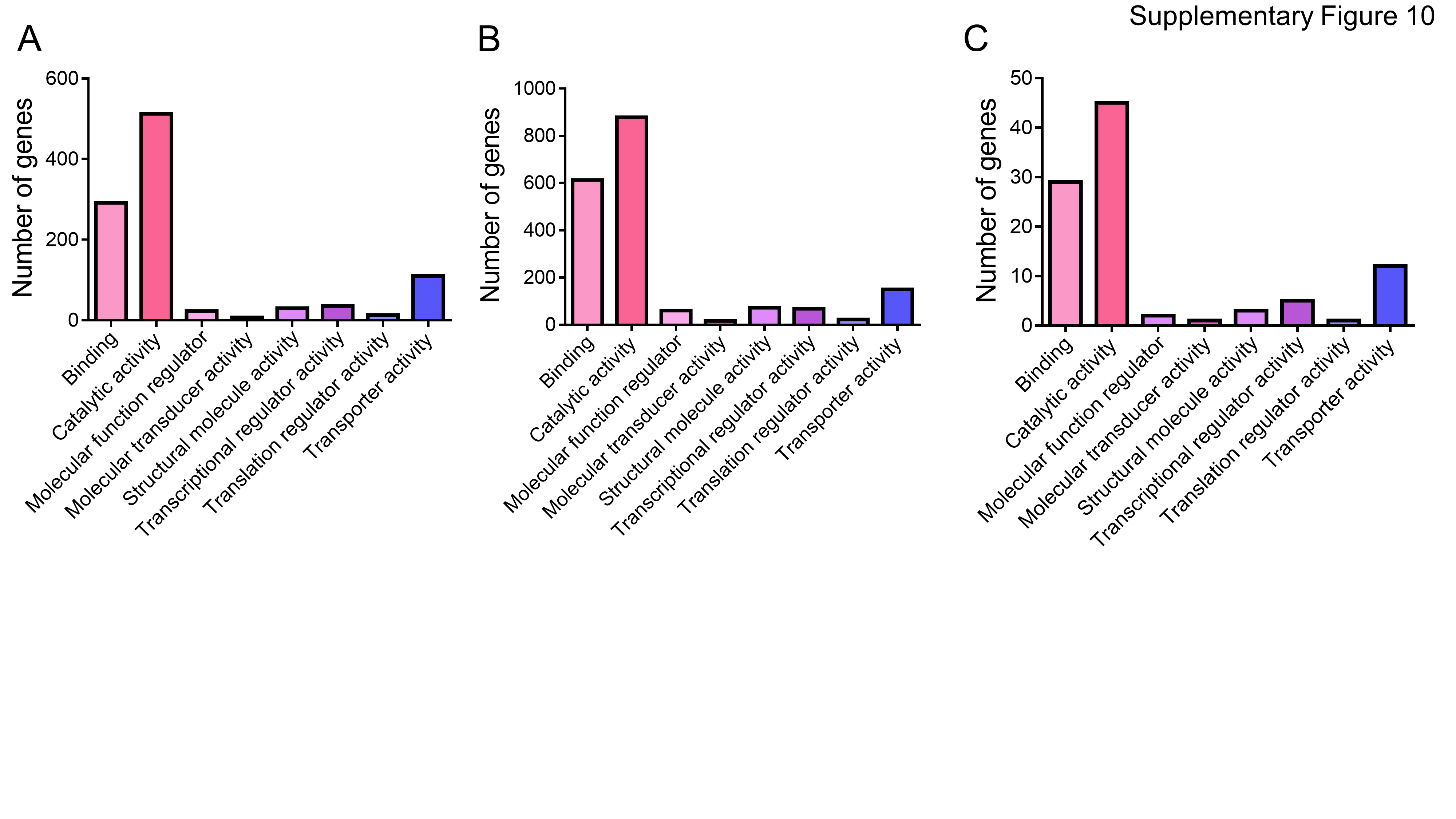

### Supplementary Figure 11

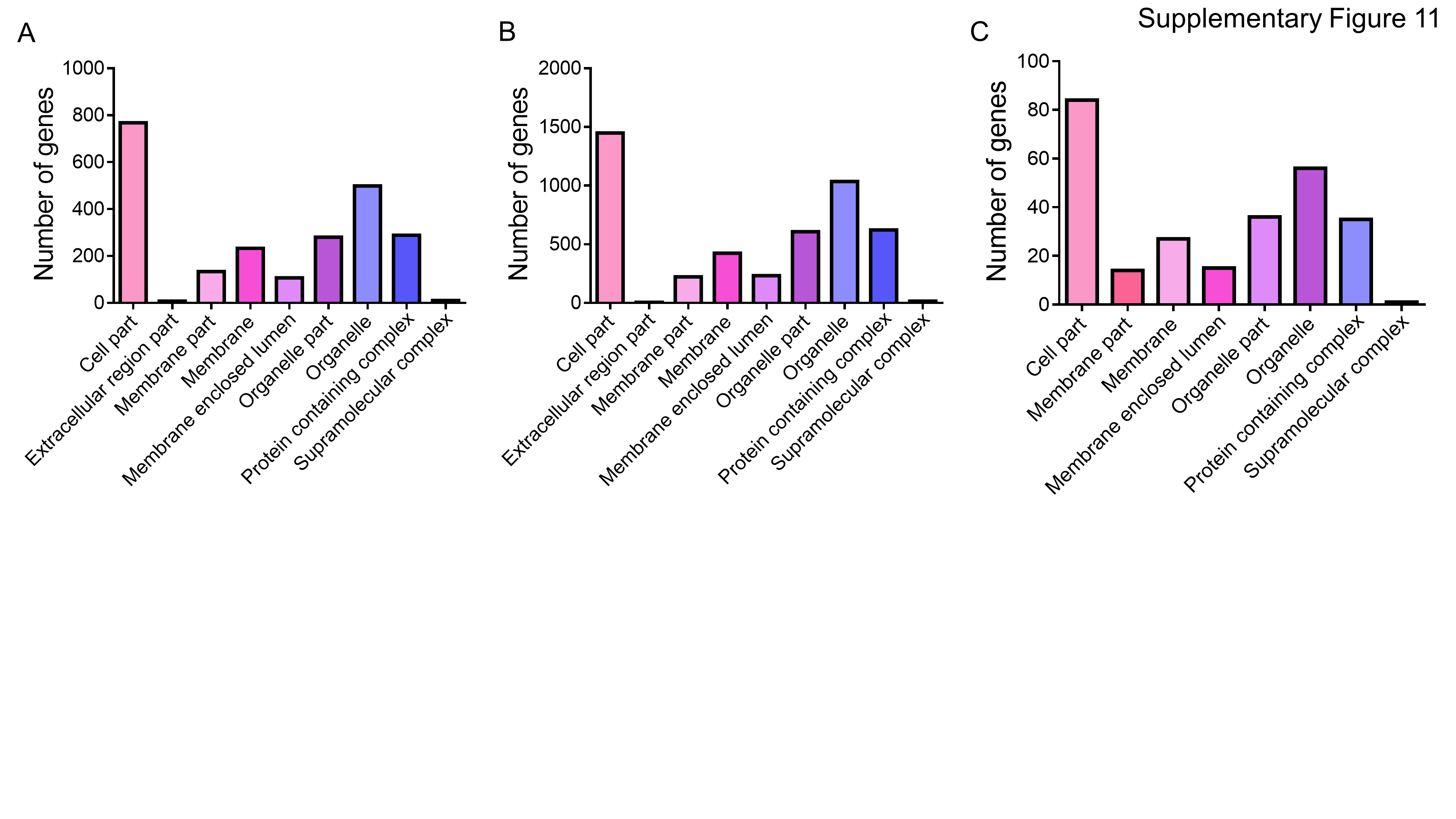

### Supplementary Figure 12

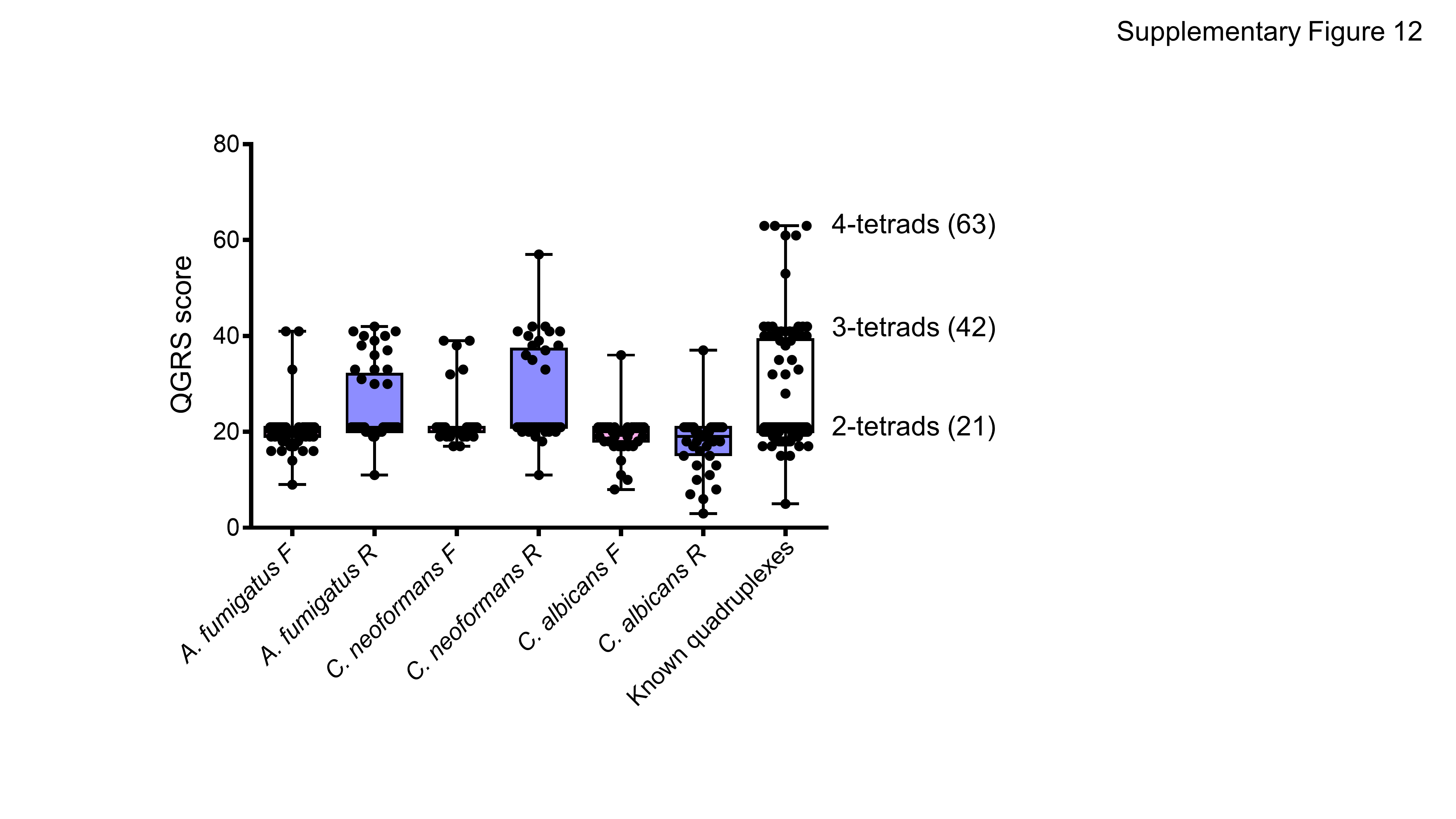
